# A different mechanism of C-type inactivation in the Kv-like KcsA mutant E71V

**DOI:** 10.1101/2021.09.22.461404

**Authors:** Ahmed Rohaim, Bram J.A. Vermeulen, Jing Li, Felix Kümmerer, Federico Napoli, Lydia Blachowicz, João Medeiros-Silva, Benoit Roux, Markus Weingarth

**Affiliations:** Department of Biochemistry and Molecular Biology, The University of Chicago, 929 E 57th Street Chicago, Illinois 60637, United States; NMR Spectroscopy, Bijvoet Center for Biomolecular Research, Department of Chemistry, Faculty of Science, Utrecht University, Padualaan 8, 3584 CH Utrecht, The Netherlands; Department of Biomolecular sciences, School of Pharmacy, The University of Mississippi, Oxford, Mississippi, United states

## Abstract

A large class of K^+^ channels display a time-dependent phenomenon called C-type inactivation whereby prolonged activation by an external stimulus leads to a non-conductive conformation of the selectivity filter. C-type inactivation is of great physiological importance particularly in voltage-activated K^+^ channels (Kv), affecting the firing patterns of neurons and shaping cardiac action potentials. While understanding the molecular basis of inactivation has a direct impact on human health, its structural basis remains unresolved. Knowledge about C-type inactivation has been largely deduced from the pH-activated bacterial K^+^ channel KcsA, whose selectivity filter under inactivating conditions adopts a constricted conformation at the level of the central glycine (TTV**<u>G</u>**YGD) that is stabilized by tightly bound water molecules. However, C-type inactivation is highly sensitive to the molecular environment surrounding the selectivity filter in the pore domain, which is different in Kv channels than in the model KcsA. In particular, a glutamic acid residue at position 71 along the pore helix in KcsA is consistently substituted by a nonpolar valine in most Kv channels, suggesting that this side chain is an important molecular determinant of function. Here, a combination of X-ray crystallography, solid-state NMR and molecular dynamics simulations of the E71V mutant of KcsA is undertaken to explore the features associated with this Kv-like construct. In both X-ray and ssNMR data, it is observed that the filter of the Kv-like KcsA mutant does not adopt the familiar constricted conformation under inactivating conditions. Rather, the filter appears to adopt a conformation that is slightly narrowed and rigidified over its entire length. No structural inactivation water molecules are present. On the other hand, molecular dynamics simulations indicate that the familiar constricted conformation can nonetheless be stably established in the mutant channel. Together, these findings suggest that the Kv-like E71V mutation in the KcsA channel may be associated with different modes of C-type inactivation, showing that distinct selectivity filter environments entail distinct C-type inactivation mechanisms.

## INTRODUCTION

Voltage-gated potassium (Kv) channels control the selective flow of potassium (K^+^) across cell membranes, which is vital for electric signalling.^1^ Kv channels are important therapeutic targets^2^ and composed of four identical subunits with six transmembrane segments. The first four segments form the voltage sensor and the two terminal segments the central pore (Figure 1a). The pore includes the selectivity filter, a conserved molecular gate that governs ion selectivity. The selectivity filter has the canonical signature sequence TVGYG conserved among all K^+^ channels^3^ and its backbone carbonyl-groups together with the threonine sidechain line up to form five K^+^ coordination sites S0-S4.^4,5^ The function of Kv channels is regulated by two distinct inactivation mechanisms: ^1,6–8^ N-type inactivation, an auto-inhibitory process in which the pore gets blocked by a molecular plug, and C-type inactivation, a regulatory process in which the filter adopts a non-conducting conformation (Figure 1b).^6^ Aberrations in C-type inactivation are closely related to neurological disorders such as episodic ataxia or the long QT syndrome^9^, which makes a structural framework of C-type inactivation in Kv channels desirable. The phenomenon of C-type inactivation was most clearly documented with functional measurements in the case of voltage-activated *Shaker* K^+^ channel from *Drosophila melanogaster* (the *Shaker* channel).^8,10–17^

**Figure 1.**
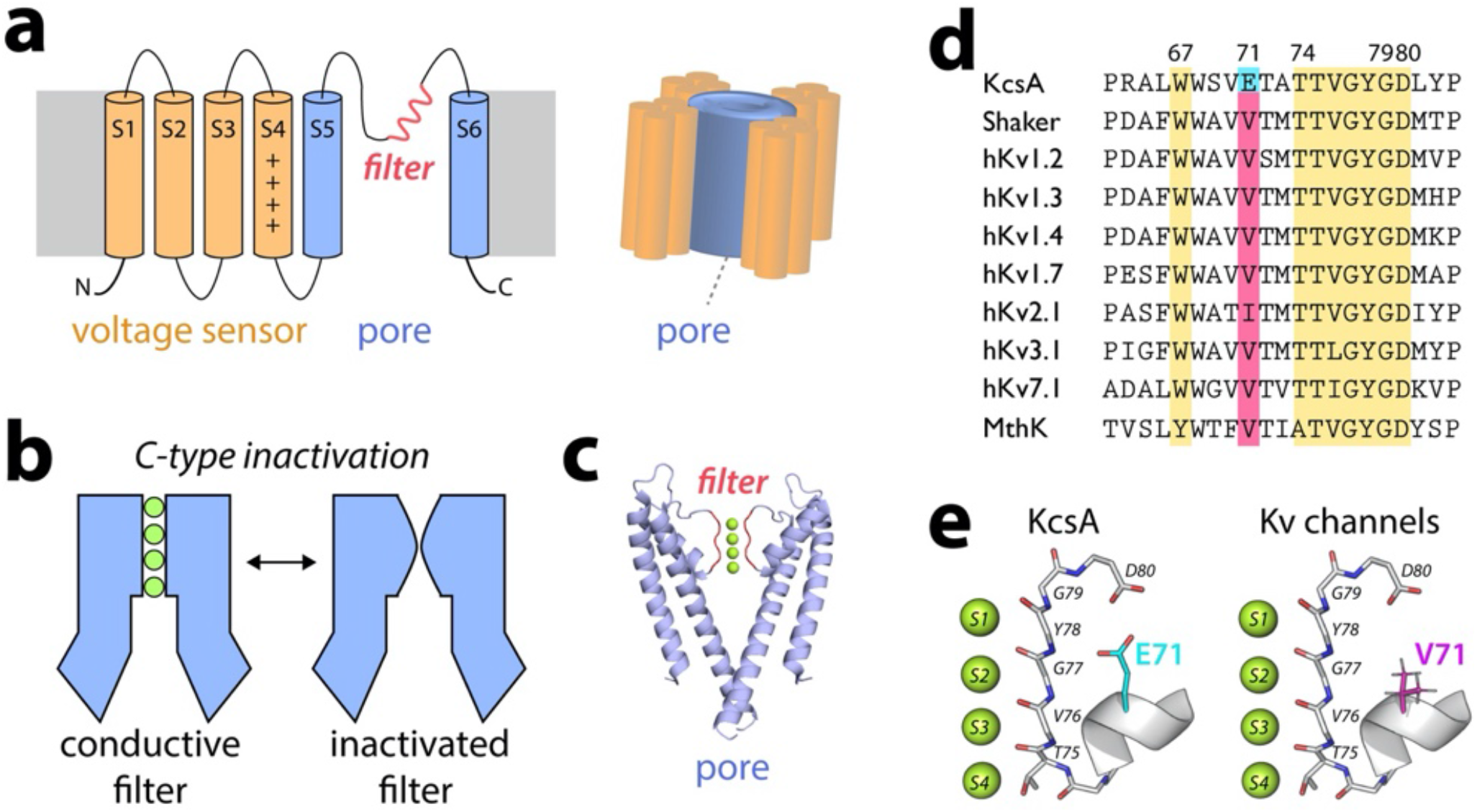
C-type inactivation and selectivity filter gating. a) Topology diagram of voltage-gated K^+^ (Kv) channels. Helices S1-S4 form the voltage-sensor, while helices S5-S6 form the pore comprising the selectivity filter. b) Slow C-type inactivation renders the selectivity filter of K^+^ channels non-conductive. c) The bacterial KcsA channel is a model for the pore of Kv channels. d,e) The selectivity filter (T74-G79 in KcsA) and residues W67, D80 are conserved while the residue corresponding to E71 in KcsA is substituted by a valine in Kv channels.

A molecular mechanism of C-type inactivation considered representative for the central pore of K^+^ channels (Figure 1c) has been established for the bacterial channel KcsA on the basis of functional measurements,^18–20^ X-ray crystallography^5,21–24^, NMR spectroscopy,^25,26^ and molecular dynamics simulations.^27–30^ Under inactivating low K^+^ conditions, the selectivity filter of KcsA sharply constricts at the level of the central glycine (TTV**<u>G</u>**YGD) caused by a rotation of the V76-G77 plane, which prevents K^+^ binding at the sites S2 and S3.^5,21^ This long-lived constricted conformation is stabilized by the binding of a few water molecules into a cavity behind the filter.^25,27^ The conductive-to-constricted transition of the selectivity filter is allosterically enhanced by the opening of the intracellular activation gate.^21,22,29^ The broad features of a KcsA-like constricted state are recapitulated in a computational study based on homology models of the pore domain of the *Shaker* channel, but several differences were noted.^31^ Clearly, C-type inactivation does not only depend on the selectivity filter signature sequence, but is exquisitely sensitive to the surrounding residues in the pore domain,^18,20,32,33^ which differ in Kv channels and KcsA (Figure 1d,e). Of particular interest, the critical glutamic acid E71^18^ along the pore helix behind the filter, known to dramatically affect inactivation kinetics of the KcsA channel^18,20,21^, is replaced by a nonpolar valine that is highly conserved in most Kv channels. This raises the question of whether C-type inactivation of Kv channels, by virtue of such a local change in the environment of the selectivity filter, could involve different or additional states than the familiar constriction observed in WT KcsA. Alternative mechanisms have been hypothesized and, despite the progress, the subject remains controversial.^34^ Putting aside the effect of voltage-sensors, the E71V mutant of KcsA offers a great opportunity to have an incisive look at the molecular environment around the selectivity filter representative to that of a Kv channel. A number of recent experimental studies challenge the canonical constricted conformation as a molecular correlate of C-type inactivation, which might not be necessarily associated with a unique selectivity filter conformation.^13,35–39^ Our knowledge of the structural basis for C-type inactivation in Kv channels still needs to be firmly established. In the following, we document a series of observations from X-ray crystallography, solid-state NMR (ssNMR), and molecular dynamics simulations showing how the inactivation process in the Kv-like KcsA mutant could be supported by a mechanism that differs from that of the WT channel (Figure 1e).

## RESULTS

### Filter narrowing in the Kv-like channel

To study the molecular mechanism of C-type inactivation in E71V KcsA, we solved X-ray structures with a closed and open intracellular gate, henceforth called closed-gate and open-gate structures (Supplementary Table 1, Supplementary Figure 1). Closed-gate structures were solved as previously described.^4,18^ To determine open-gate E71V KcsA structures, the lower ends of the two long transmembrane helices (TM1 and TM2) were cross-linked via disulfide bridges formed between residue C28 of one chain and C118 of the neighboring chain, as previously described for KcsA^23^ (Supplementary Figure 1a). Well-defined electron density maps were obtained for the transmembrane domain and the pore-loop region including the selectivity filter. Closed-gate E71V KcsA structures have resolutions in the range of 2.7-2.9 Å, whereas the resolutions of the open-gate structures are in the range of 3.0-3.3 Å.

The closed-gate E71V KcsA structure solved with 5 mM K^+^ (pdb 7MHR) closely matched the closed-conductive structure of WT KcsA (pdb 1K4C), with an overall rmsd (root-mean-square deviation) of 0.3 Å (Supplementary Figure 1b). The conformation of the closed inner gate of E71V is identical to that in WT KcsA, and the Cα-Cα distance between Thr112 of the TM2 helix is 12 Å for both channels (Supplementary Figure 1a,b). The E71V selectivity filter is similar to the WT KcsA (rmsd^T74-D80^ = 0.20 Å) with the exception of a widening of the pore outermost S1 site (Figure 2 a,b), with the Cα-Cα distance between opposing G79 residues being 8.7 Å in E71V and 7.9 Å in WT KcsA.

**Figure 2.**
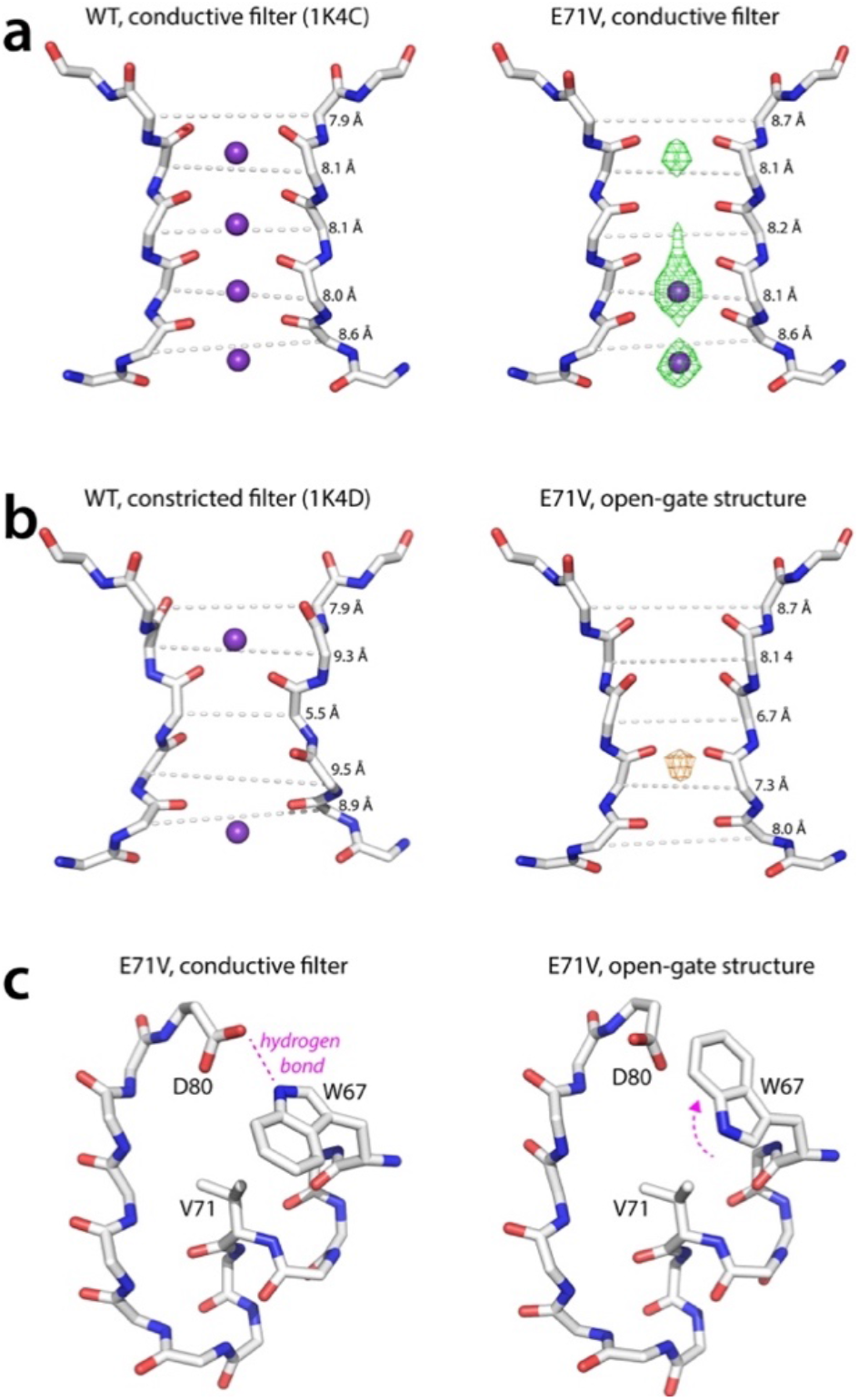
Closed-gate and open-gate X-ray structures of WT and E71V KcsA. The selectivity filter of the a) closed-gate and b) opengate X-ray structures of WT and E71V KcsA. While the WT filter shows a sharp constriction at G77, the E71V KcsA filter shows no constriction but a pore narrowing at sites S2-S4. The Cα-Cα distances between residues of opposing protomers are shown. The closed-state and open-state E71V KcsA structures were obtained with 5 mM and 0 mM K^+^, respectively. The green mesh shows the difference density map, the orange mesh the anomalous density. c) The W67 sidechain flips in the E71V open-state structure, breaking the hydrogen bond with the D80 sidechain that is important for inactivation in Kv channels.^12,19,40,41^

The open-gate E71V structure (pdb 7MK6) was solved at low K^+^ (0 mK^+^/150 mM NaCl), which is an inactivating condition for KcsA.^5,21,22^ The crystal structure globally resembles the previously described open-gate WT KcsA structures^22,23,42^ (Supplementary Figure 1b). The opening extent of the inner gate is similar to an opengate structure of WT KcsA in which TM1 and TM2 were cross-linked,^42^ with a Cα-Cα distance at position Thr112 roughly 22.5 Å for both structures (Supplementary Figure 1a). Strikingly, the E71V open-gate selectivity filter does not adopt the canonical constricted conformation observed in low K^+^ WT KcsA (pdb 1K4D) (rmsd_T74-D80_ = 0.67 Å) and open-gate WT KcsA structures (pdb 3F7V) (rmsd_T74-D80_ = 0.78 Å). As shown in Figure 2b, the filter in the open-gate E71V structure is not constricted at the level of V76-G77. Rather, the entire filter-pore appears to be narrower in the open-gate structure compared to the close-gate structure. The narrowing is most pronounced at the S2 site (Cα-Cα distance of opposing G77 residues reduced by 1.5 Å), and still sizeable at the S4 site (Cα-Cα distance of opposing T75 residues reduced by 0.6 Å), while the S1 site and the pore mouth (Y78-D80) are virtually unaffected. These structural features depart from the constricted filter of WT KcsA, in which the S1 and S4 sites widen as a consequence of the sharp constriction at the level of the central glycine residue. The pore-narrowing in E71V is accompanied by only small structural changes compared to the conductive filter (rmsd_T74-D80_ = 0.42 Å), and dihedral angles of filter residues alter by 20-25°.

Interestingly, upon opening of the inner gate of E71V, the W67 sidechain adopts an alternative rotamer conformation, breaking the hydrogen bond with D80 (Figure 2c). Several studies reported on the importance of the W67 – D80 hydrogen bond for inactivation in Kv channels^12,19,40,41^, and breaking this interaction enhances inactivation to different degrees. In *Shaker*, mutation of the corresponding residue W434 to a F results in near-constitutively inactivated channels,^40^ whereas the corresponding mutation of W366 in Kv1.2 accelerates inactivation^19^, while the mutation of D317N (D80 in KcsA) in Kv7.1 abolishes ion conductance ^41^. Presumably, the conformational change of W67 is allosterically induced by the opening of the very distant inner gate. This is supported by the observation that the F103 sidechain, an important determinant of the gate-filter allosteric coupling in KcsA^22^, also changes its conformation upon channel opening and moves closer to T74 (Supplementary Figure 1c).

Next, we compared inactivation gating in full-length E71V and WT KcsA using ssNMR, which enables high-resolution studies of proteins in a fluid membrane. High-quality ssNMR data were acquired with [^13^C,^15^N]-labelled inversely fractionally deuterated^43^ channels under high K^+^/neutral pH conditions (100 mM K^+^/pH7) at which channels are closed with a conductive filter (closed-conductive), and under low K^+^/acid pH conditions (0 mM K^+^/pH3-4) at which channels are open with an inactivated filter (open-inactivated)^20^ (Figure 3a,b). Using our previous chemical shift assignments of closed-conductive WT KcsA as reference,^44^ backbone assignments were obtained for all states using 3D (CANH, CONH, CAcoNH, COcaNH) and 2D CC data (Supplementary Figures 2-5).

**Figure 3.**
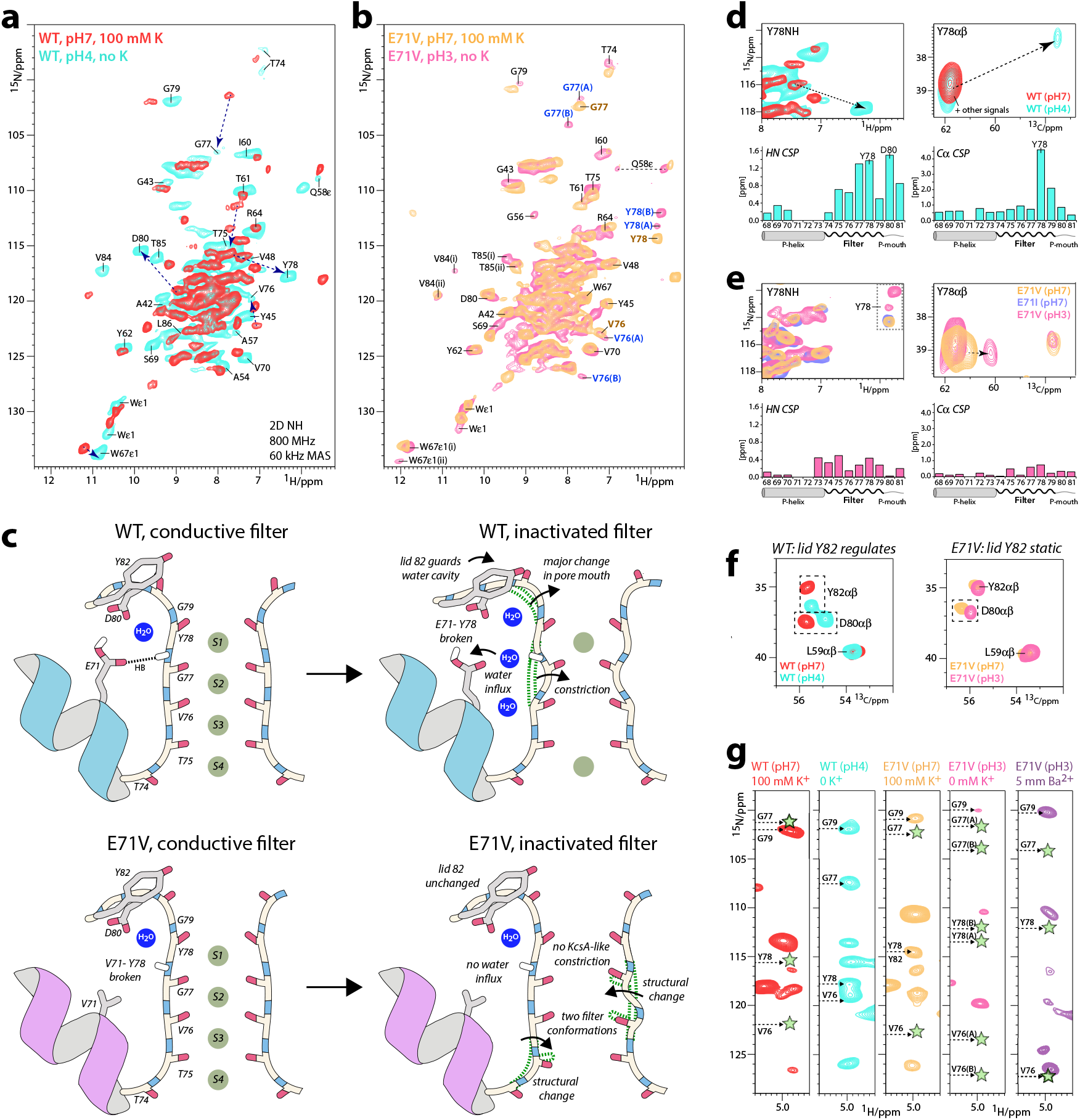
Comparing inactivation gating by ssNMR. a) Superposition of 2D NH ssNMR spectra of a) closed-conductive (red) and open-inactivated (cyan) WT KcsA channels acquired in membranes. Arrows indicate major signal shifts of key residues. b) 2D NH spectra of E71V KcsA in closed-conductive (yellow) and open-inactivated (pink) states. The E71V KcsA filter shows two conformations *A* and *B*, highlighted with blue labels, under inactivation conditions. V76, G77, and Y78 of the canonical conductive filter (1K4C) are labelled in yellow for clarity. c) Schematics of the changes that accompany inactivation in WT and E71V. The inactivated WT filter constricts at sites S2 and S3, accompanied by a rupture of the E71 – Y78 interaction and the influx of water into a cavity behind the filter. NMR data also reveal major structural changes at the S1 site and the pore mouth. The inactivated E71V filter does not constrict like KcsA, does not feature structural water in the cavity, and shows two conformations. d,e) Upper panel: 2D NH and 2D CC spectra of the WT filter showing large chemical shift perturbations (CSPs) of key residue Y78 upon constriction. Lower panel: HN and Cα CSPs of the open-inactivated state in reference to the closed-conductive state. Source data are provided as a Source Data file. e) Same as d) for E71V, which shows only small CSPs upon inactivation. f) Residue Y82 guards the access to the water-cavity. 2D CC spectra show that Y82 undergoes a conformational change upon constriction of the WT filter but is static during E71 inactivation. g) 2D N(H)H ssNMR spectra to probe structural water behind the conductive (red) and constricted (cyan) WT filter, the conductive (yellow) and inactivated (pink) E71V filter, and the barium-bound E71V filter (magenta). Green stars mark the absence of structural water.

First, we used ssNMR to show that the WT KcsA selectivity filter and its environment undergo large conformational changes upon inactivation (Figure 3c,d). As previously reported,^26^ we observed a strong chemical shift perturbation (CSP) for the carbonyl of V76 (+2.99 ppm relative to the conductive WT filter) (Supplementary Figures 4,5), which is the signature for the constricted filter of WT KcsA.^26^ In addition, we detected large CSPs for residues ^1^H- and ^13^Cα-CSPs for Y78-L81 that report marked conformational changes at the outermost S1 site at the pore mouth (Figure 3c,d and Supplementary Figure 6). As shown in the X-ray structure of WT KcsA (1K4D)^5^, these changes are caused by the loss of the E71 – Y78 interaction behind the constricted filter.

Next, we used ssNMR to study selectivity filter gating in the E71V pore. The chemical shifts of the conductive filters of E71V and WT KcsA closely resemble each other except for stark local CSPs of residues W67, Y78 and D80 (Supplementary Figure 7) that are, presumably, due to the nonpolar valine residue at position of residue 71. The valine cannot maintain an interaction with Y78/D80 as in WT KcsA, leading to major conformational differences at pore mouth and S1 site in E71V compared to WT KcsA. This reasoning is consistent the minor CSPs that we observed between conductive and inactivated selectivity filters of E71V KcsA. Since there is no interaction between V71 of the pore helix and Y78 in E71V, the pore mouth and S1 site remain largely unaffected during C-type inactivation (Figure 3c,e). This appears to be consistent with the minor changes at the S1 site and pore mouth between closed-gate and open-gate E71V in X-ray structures. Note that the ssNMR observations for E71V KcsA are reproduced if residue E71 is replaced by an isoleucine,^44^ encountered in some Kv channels (Figure 3e).

The entire E71V selectivity filter shows signal splittings (Figure 3b), which means that the pore adopts either one asymmetric or two different states under inactivating conditions. None of the two E71V filter conformations, henceforth called conformations *A* and *B*, appear to adopt the constricted conformation that WT KcsA features at the same experimental conditions.^5,26^ This is evidenced by strong signal difference of the V76CO signature signal (+3.10 ppm in conformation *A*, +3.60 ppm in conformation *B*) compared to the constricted WT filter^26^ (Supplementary Figures 3-5). Conformation *A* of the inactivated E71V filter shows marked CSPs at the S3 and S4 sites for T75CO (−1.2 ppm in reference to the E71V conductive filter) and V76Cα (+0.8 ppm), but otherwise resembles the conductive conformation (Figure 5h). Conformation *B* shows a major CSP for G77CO (+1.95 ppm) at the filter S2/S3 sites. The perturbations observed at T75, V76, and especially G77 appear to be consistent with the open-gate X-ray structure of E71V, which shows a pore narrowing for the S2 – S4 sites, which is most pronounced at G77.

The W67 sidechain, critical for inactivation in Kv channels,^12,19,40,41^ also shows two conformations in ssNMR spectra of E71V under inactivating conditions (Figure 3b). Signal splittings are also observed for V84 and T85, right above the W67 sidechain in the adjacent protomer, that tightly interact^45^ with the E51 sidechain that plays an important role in C-type inactivation and the coupling between pore and voltage-sensor in *Shaker*.^46^

These observations match with structural changes for E51, V84, and T85 observed in the open-gate E71V X-ray structure (Supplementary Figure 8). We also detected a second conformation for T74 that is part of the allosteric network that couples the inner gate to the filter.^22^ Together, the ssNMR data strongly suggest that the conformations and detailed interactions of W67 and V84/T85 are modulated allosterically. Note that because the asymmetric unit of the E71V KcsA crystals contains only one monomer, the generated tetrameric biological unit is necessarily four-fold symmetric. A conformation departing from the four-fold symmetric tetramer would only be observable by X-ray if the channel crystallized in an asymmetric unit containing more than a single monomer.^47^

The influx of water into the cavity behind the filter, guarded by lid residue Y82,^23,25,27^ is crucial for the long-lived inactivation of WT KcsA.^25,27^ Using 2D ^15^N(^1^H)^1^H ssNMR experiments^48^ that monitor the proximity of rigid water to backbone amino-protons, we could confirm the presence of several water molecules behind residues V76-G79 of the WT KcsA constricted filter, while water is confined to G79 behind the conductive filter (Figure 3g). Moreover, upon inactivation, we observed a marked signal change (+1.29 ^13^Cβ ppm) of the Y82 sidechain, consistent with a conformational change of the lid Y82^23,25,27^. Conversely, no conformational change of Y82 was observed in E71V KcsA under conditions promoting inactivation (Figure 3f). We therefore hypothesized that water plays a less prominent role behind the filter of Kv channels. Indeed, ssNMR experiments show that the cavity behind the inactivated E71V filter is devoid of rigid water molecules, in line with the hydrophobic nature of the valine 71 sidechain (Figure 3g). This appears to be another major difference to inactivation in KcsA and suggests that inactivating water is not a universal feature of C-type inactivation, which aligns with observations for C-type gating in the human K_2P_2.1 (TREK-1) channel^49^ and the MthK channel.^50^ In line with ssNMR, no water molecule was observed in the cavity behind the selectivity filter of the open-gate E71V X-ray structure. Although our ssNMR setup excludes the presence of rigid water molecules with resistance times in the microsecond range (or longer), transient visits of fast exchanging water behind the filter cannot be ruled out.

### Effect of barium

We used barium to further highlight the different conformations adopted by the E71V channel under inactivating conditions compared to the WT channel. Ba^2+^ can serve as a very useful biophysical probe of the selectivity filter. The cation has a similar crystal radius compared to K^+^, yet it tightly binds to the filter and blocks ion conduction.^42,51–55^

In WT KcsA, Ba^2+^ primarily binds at the S4 site in both closed-gate (pdb 2ITD^54^ and 6W0I^42^) and open-gate (pdb 6W0D^42^) X-ray structures (Figure 4a,b). The selectivity filter of Ba^2+^-bound closed-gate WT KcsA is constricted with a G77-G77 distance of 5.4 Å, similar to 1K4D. The Ba^2+^-bound open-gate WT KcsA corresponds to a semiconstricted filter with a G77-G77 distance of 6.0 Å with some structural deviations that occur at the T75 sidechain.

**Figure 4.**
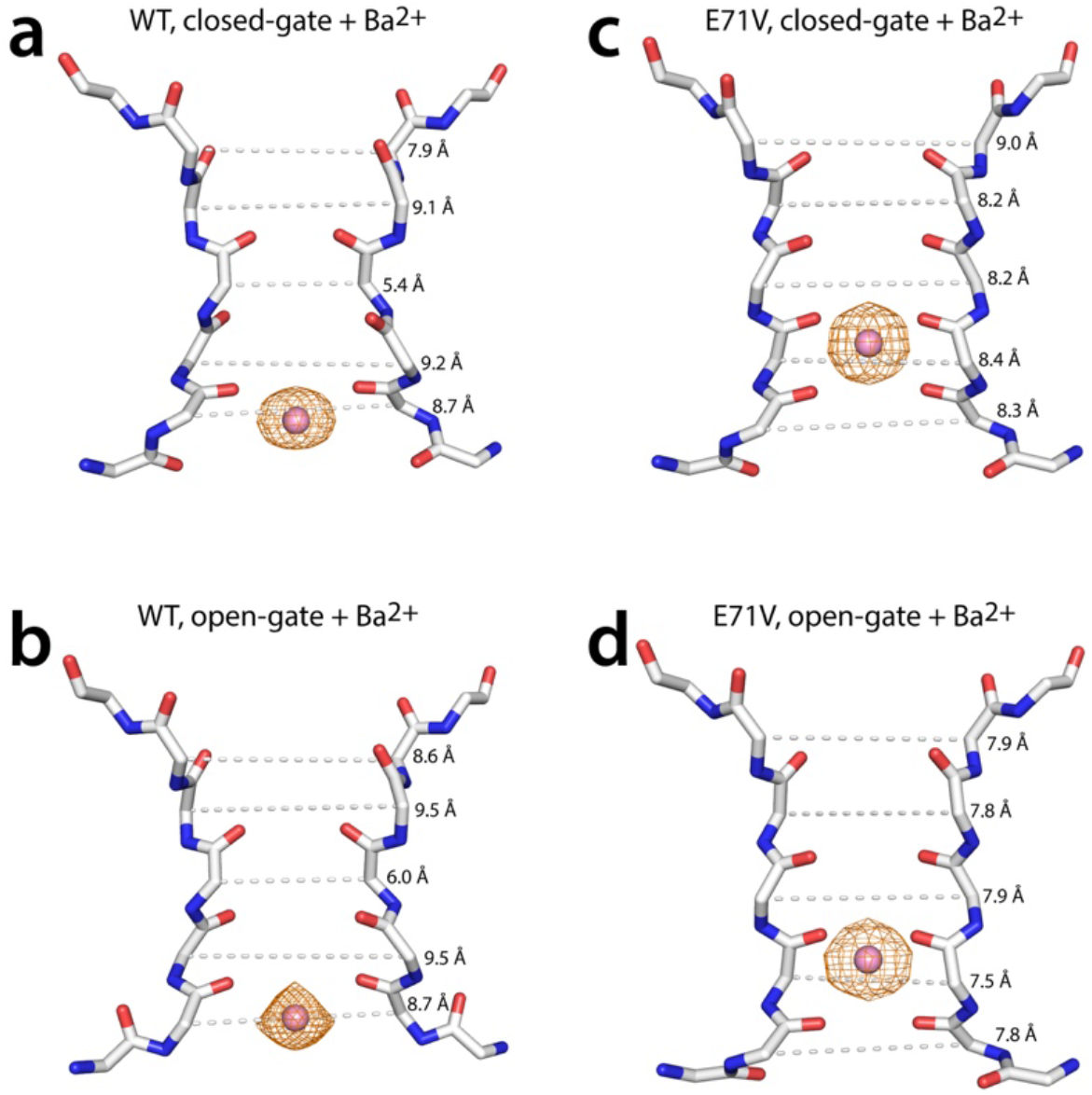
X-ray structures of barium-bound WT and E71V KcsA. a) Closed-gate and b) open-gate structures of WT KcsA with Ba^2+^ bound at the S4 position of the selectivity filter. The filter is sharply constructed at the level of G77 in both the closed-gate and in the opengate structures. c) Closed-gate and d) open-gate structures of E71V KcsA with Ba^2+^ bound at the S3 position of the selectivity filter. The filter is conductive in the closed-gate structure and narrows in the open-gate structure. Cα-Cα distances between like-residues in opposing protomers are indicated.

**Figure 5.**
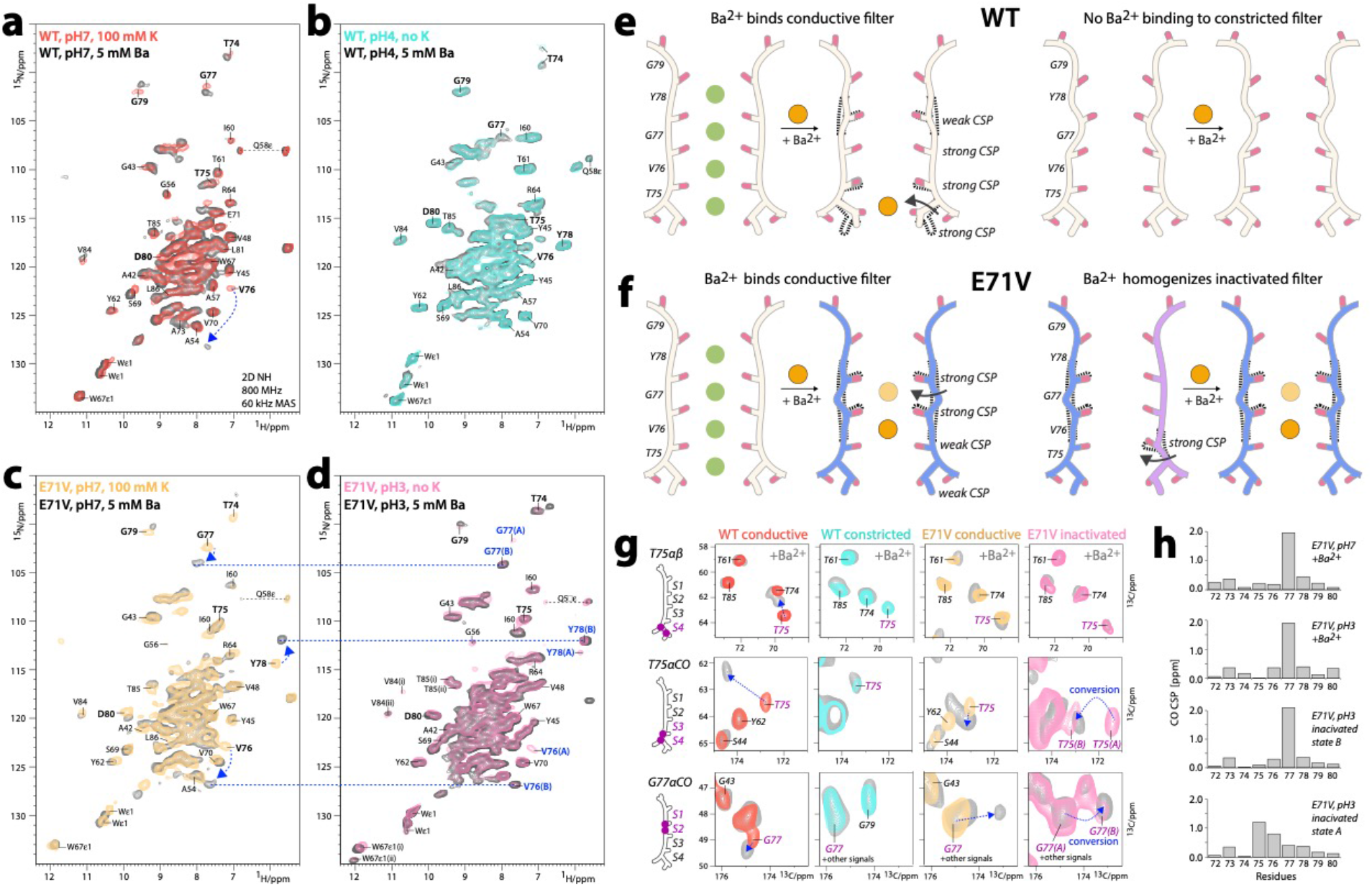
ssNMR barium-binding studies. 2D NH spectra of a) closed-conductive WT KcsA (red), b) open-inactivated WT (cyan), c) closed-conductive E71V KcsA (yellow), and d) open-inactivated E71V (pink). Superimposed (in grey) are 2D NH spectra after equilibration with 5 mM Ba^2+^ solution of the same pH as the original sample. Major Ba^2+^-caused signal changes are indicated by arrows. The two conformations *A* and *B* of the E71V filter under inactivating conditions are highlighted by blue labels. Blue lines show that Ba^2+^ binds the E71V filter with a conformation similar to conformation *B* at neutral and acidic pH. e,f) Schematics of Ba^2+^ binding to the WT and E71V selectivity filters. Barium binds at the S4 site of the conductive, however not the constricted WT filter. Barium binds the conductive and the inactivated E71V filter at the S3 or S2 site. g) 2D CC spectra show the effect of Barium binding to key residues T75 and G77 of the S2-S4 filter sites. h) Backbone carbonyl CSPs in reference to the conductive E71V filter show that the Ba^2+^-bound states of E71V at pH7 and pH3 closely resemble state *B* of the inactivated E71V filter. Source data are provided as a Source Data file.

To compare the effect of the E71V mutation on the selectivity filter, the crystal structures of closed and open-gate E71V KcsA were determined in the presence of Ba^2+^ and in the absence of K^+^ (Figure 4). In contrast to the WT KcsA, Ba^2+^ predominantly binds at the S3 site in both E71V structures, which is confirmed by the anomalous x-ray diffraction as shown in Figure 4. In the Ba^2+^-bound closed-gate E71V structure, the selectivity filter is similar to the conductive state without Ba^2+^ (rmsd_T74-D80_ = 0.15 Å) with a G77-G77 distance of 8.2 Å (Figure 2a, 4c). On the other hand, the open-gate Ba^2+^-bound E71V structure is narrowed compared to the closed-gate Ba^2+^-structure (Figure 4d). This aligns with the selectivity filter narrowing observed for the open-gate E71V KcsA crystal structure in absence of Ba^2+^ (Figure 2b). In addition, the open-gate E71V structures, with and without Ba^2+^, show the same conformational change of the W67 sidechain if compared to WT KcsA.

Next, we equilibrated E71V and WT KcsA proteoliposomes with 5 mM Ba^2+^ at neutral (pH7) and acidic (pH3-4) conditions to study Ba^2+^ binding to the closed-conductive and open-inactivated states by ssNMR. For WT KcsA at pH7, we observed major CSPs for the T75 backbone and sidechain and the V76 backbone (Figure 5a, e, g) which implies that Ba^2+^ binds to the S4 site, in line with X-ray studies.^42,54^ The signature CSP of V76CO, which reports on the constriction^26^, is −2.99 CO ^13^C ppm for the constricted filter (compared to the conductive WT filter) and only −2.03 CO ^13^C ppm in the Ba^2+^-bound filter (Supplementary Figures 3,4). This suggests a semi-constricted Ba^2+^-bound WT filter at pH7, in line with X-ray data^42,54^. We did not observe signal changes at acidic conditions, which implies that Ba^2+^ did not bind to the constricted filter. This departs from X-ray data that observe Ba^2+^ at the S4 site of the constricted filter, however, NMR and X-ray agree that open-gate WT KcsA adopts a fully constricted filter in the presence of Ba^2+^.

In contrast to WT, Ba^2+^ binds to the E71V filter at neutral and acidic conditions in ssNMR experiments. The chemical shifts for the Ba^2+^-bound E71V filter are the same at both conditions and only one filter conformation is observed (Figure 5c, d, f, g) that is not constricted (V76CO = 179.82 ppm compared to 176.1 ppm in the constricted WT filter). In E71V, Ba^2+^ binding causes a major CSP for G77CO (+1.95 ppm relative to the conductive E71V filter) and V76NH (+1.01 ppm) at the filter S2/S3 sites (Figure 3g,h), but only minor CSPs at S4 (T75CO = +0.21; T75a CSPs = +0.45 ^13^C ppm). Hence, ssNMR data confirm the X-ray data that Ba^2+^ binds in the middle of the E71V selectivity filter (Figure 3f), which again contrasts with WT KcsA. Surprisingly, the signals of the Ba^2+^-bound E71V filter conformation closely match to filter conformation *B* that we observed at inactivating conditions (Figure 3c, d), with the same characteristic perturbation at G77CO (Figure 3g, h).

While filter residues T75-D80 adopt a single, defined state in the presence of Ba^2+^, we detected the same peak-splitting for the network T74, W67 sidechain, V84, and T85 in Ba^2+^-bound open-gate E71V that we observed at inactivating conditions without Ba^2+^. This strongly corroborates that the break of the W67 – D80 interaction is allosterically caused by the opening of the lower gate and not caused by a change in the filter conformation, because peak splittings are not observed for Ba^2+^-bound closed-gate E71V, and the filter conformation is the same for both Ba^2+^-bound channels.

### Filter rigidification upon inactivation

We probed the impact of the E71V mutation on the channel dynamics in reference to the gating mode with ssNMR relaxation, which is an ideal approach to study the internal dynamics of proteins in membranes^44^ (Figure 6a,b). We measured the ^15^N rotating-frame relaxation (*R*_1rho_) that is dominated by motion in the microsecond range.^56^ As we recently showed^44^, the selectivity filter is the most dynamic membrane-embedded region in closed-conductive WT KcsA. Unexpectedly, the E71 to V substitution enhanced the dynamics of the entire closed-conductive channel, with a pronounced local surge in dynamics at V76 in middle of the filter, in direct vicinity to the V71 sidechain (Figure 6 d, e).

**Figure 6.**
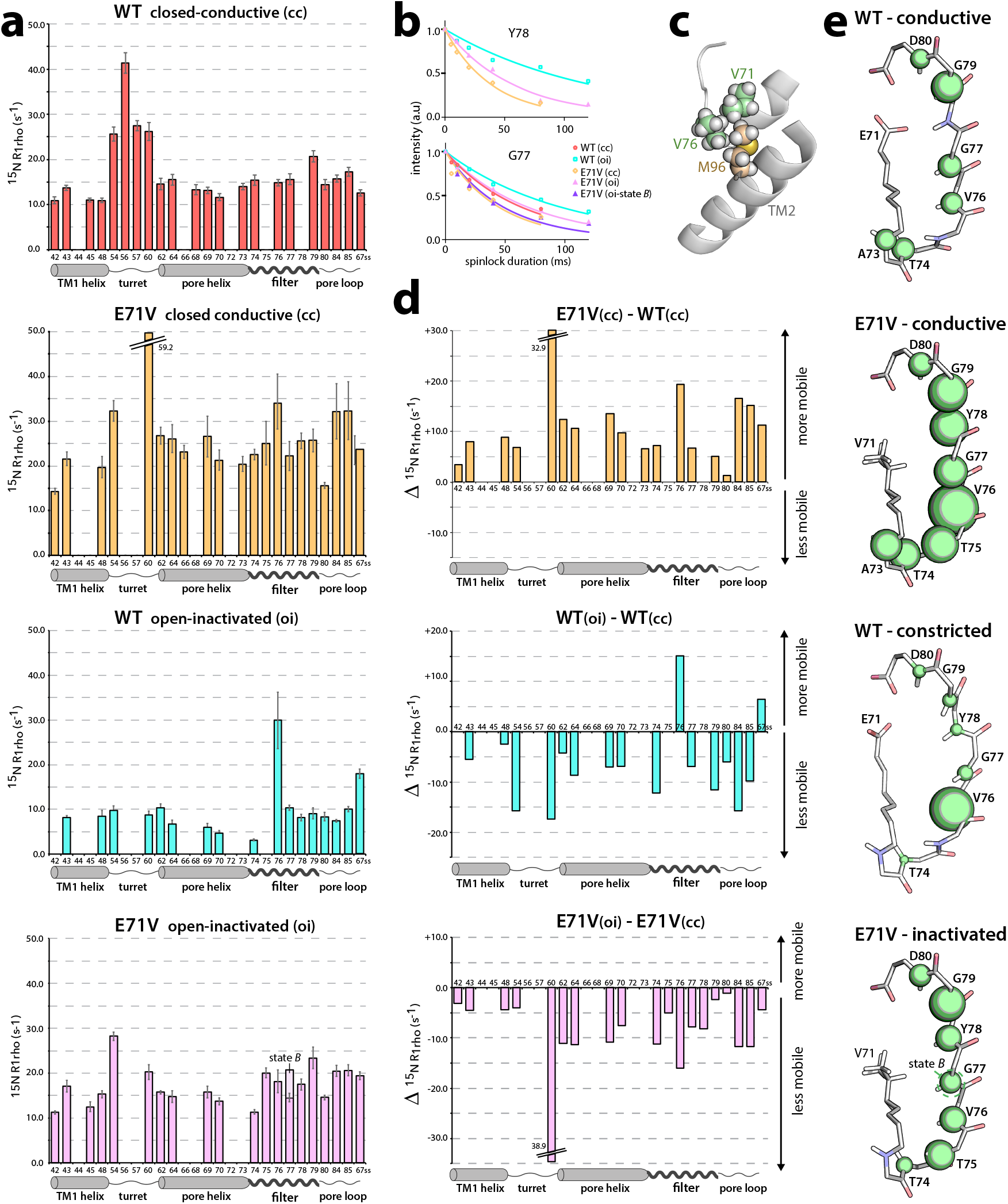
Dynamics measurements in different gating modes. a) ^15^N rotating-frame ssNMR relaxation rates (*R*_1rho_) that report on slow molecular motions in closed-conductive (cc) and open-inactivated (oi) WT KcsA (red, cyan) and E71V (yellow, pink), measured at 700 MHz and 60 kHz MAS. The error bars show the standard error of the fit. Source data are provided as a Source Data file. b) *R*_1rho_ signal decay curves for selected filter residues. Symbols mark data points and lines represent best fits. c) The valine behind the E71V filter is in direct contact with the hydrophobic V76 and M96 sidechains. d) Plots of the differences in the dynamics between the closed-conductive (cc) states of E71V and WT, and for each WT and E71V the dynamics differences between their open-inactivated (oi) and closed-conductive (cc) states. e) Illustration of the site-resolved selectivity filter dynamics. The size of the green spheres represents the *R*_1rho_ relaxation rates.

Steric clashes between the V71, V76, and M96 sidechains could possibly cause this increase in mobility (Figure 6c), analogous to the increase in dynamics observed in the E71I mutant caused by clashes between the I71 and W67 sidechains.^44^

Both channels are globally more rigid at inactivating conditions (Figure 6d,e), presumably due to the opening of the lower gate, in line with EPR data that showed a stiffening of open-gate WT.^57^ In WT, we observed a distinct local increase in the mobility of V76 at the S2/S3 sites. This means that the constriction is a flexible structural element, which aligns with computational results^28^. In contrast to WT, all residues of the S2 and S3 sites in E71V rigidify upon inactivation. Differences are also visible for the pore mouth and the S1 site that rigidity in WT upon inactivation, but are unaffected in E71V, which aligns with the absence of conformational changes at the E71V KcsA pore mouth. These data show that not only the filter structure, but also the filter dynamics differ between E71V and WT KcsA. We also observed a significant difference for G77 between states *A* (~ 14 ms^-1^ *R*_1rho_) and *B* (~20 ms^-1^ *R*_1rho_) of E71V (Figure 6a). This agrees with G77 being the residue with the most pronounced chemical shift difference between *A* and *B* states (Figure 5h).

### Outlook from molecular dynamics

To expand our perspective of the conformations accessible to the E71V pore, 0.5 μs molecular dynamics (MD) simulations were carried out to examine the effect of various factors. The crystal structures reported in this study were used as starting models to carry out unbiased MD simulations. In addition to the E71V mutation, we probed the effect of different ions in the selectivity filter. In the E71V open-gate, K^+^ ions were located at the sites S1 and S4 with one water molecule at S2 or S3. The K^+^ in S1 maintained its position, however the K^+^ in S4 was delocalized to S3 after 60 ns and stayed there for the remainder of the simulation. This ion movement was associated with a slight dilation in the middle of the selectivity filter. The starting model was pre-constricted with a Cα-Cα cross-subunit distances at G77 of 6.8 Å, which is used to monitor the width of the selectivity filter. The selectivity filter remained pre-constricted for 60 ns, after which the G77 Cα-Cα cross-subunit distances widened to ~8 Å to resemble the canonical conductive conformation (1K4C) stable until the end of the simulation (Figure 7a).

**Figure 7.**
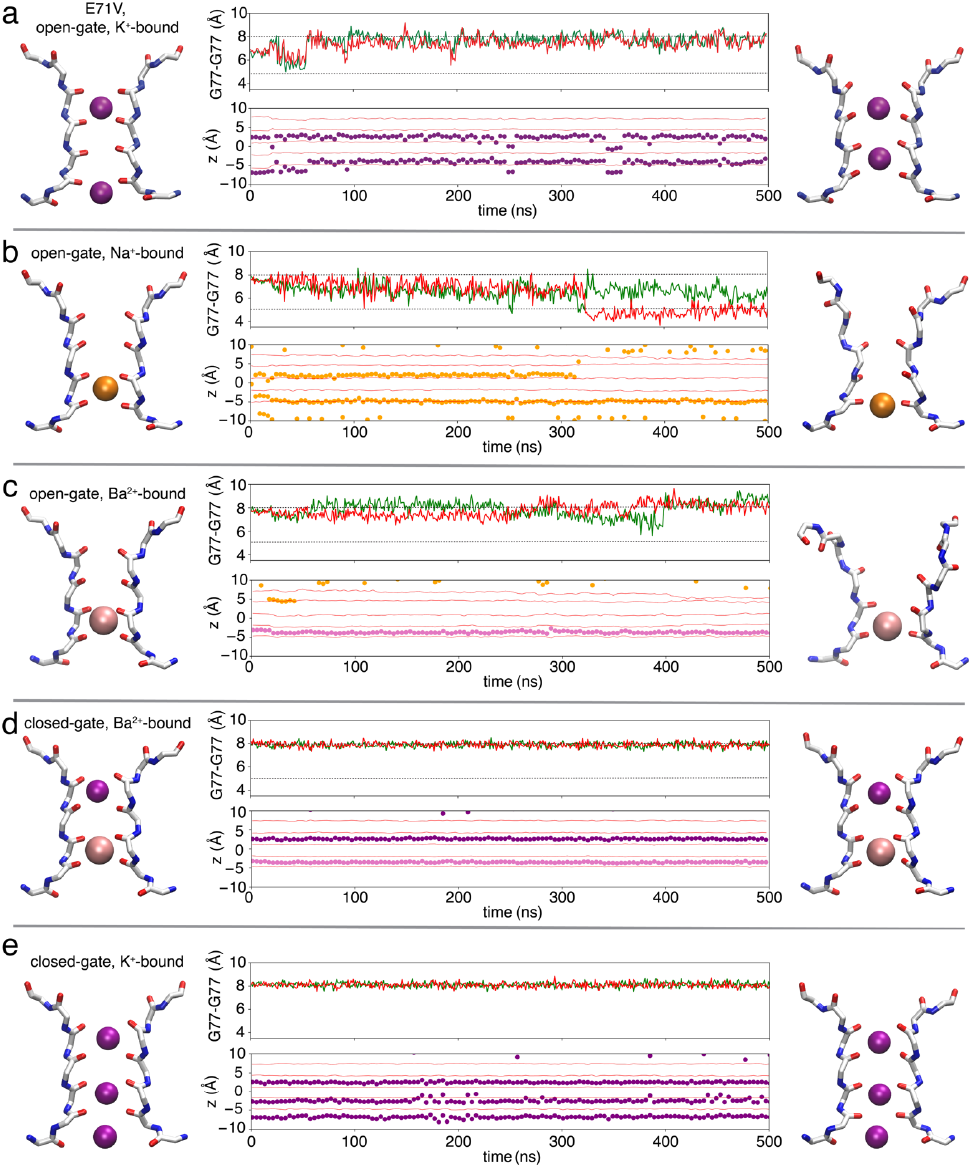
MD simulations for the structures of the KcsA E71V mutant in different conditions. (a) open-gate and K^+^ bound, (b) opengate and Na^+^ bound, (c) open-gate and Ba^2+^ bound, (d) closed-gate and Ba^2+^ bound, and (e) closed-gate and K^+^ bound. The first and last snapshots of the selectivity filter for each MD trajectory are respectively shown (right and left panel) in licorice representation for backbone and in van der Waals representations for K^+^ (purple), Na^+^ (orange), and Ba^2+^ (pink) ions. In the middle panel is the time series of the cross-subunit distance (r) between the Cα atoms of G77 of diagonally opposed monomers A and C (green), and B and D (red). Two levels representing conductive and constricted states are illustrated in black dashed line. The z-coordinate of ions in the selectivity filter are shown as solid dots (with the same colors as the conformation images). The z-coordinates of carbonyl oxygens of G79, Y78, G77, V76, and T75 are respectively shown in red lines to indicate the position of ion binding sites.

When K^+^ is substituted by Na^+^ of open-gate E71V, the sites S1-S4 of the selectivity filter were occupied interchangeably by Na^+^. After 320 ns, the filter adopted a constricted conformation with one Na^+^ still bound at the site S4 (Figure 7b). In contrast to the simulation of the open-gate E71V with K^+^ ions, the selectivity filter gradually converted from a canonical conductive to a constricted filter when K^+^ is substituted by Na^+^, similar to the constricted X-ray structure of KcsA previously observed at low K^+^ (1K4D).^5^ The G77 Cα-Cα cross-subunit distances decreased from ~ 8 Å to ~5.4 Å at 320 ns, which coincides with the unbinding of a Na^+^ ion from site S1 (Figure 7b). Remarkably, unlike the constricted form of WT KcsA, the selectivity filter of the E71V mutant constricts in an asymmetric fashion. This is shown by the green and red traces illustrating the cross-subunit distances from the opposite pairs of the monomers (Figure 7b). Additional studies reported similar asymmetrically constricted conformation in other K^+^ channels.^58^ A previous computational study of the pore domain of *Shaker* also provides indications of a structural heterogeneity associated with the constricted state of E71V;^31^ the free energy minimum in the 2D-PMF is an elongated basin covering both symmetrical and asymmetrical conformations (see Figures 6 and S3 of ref. ^31^).

We report crystal structures with Ba^2+^ binding at S3 of the selectivity filter of E71V KcsA in the open and closed gate E71V KcsA, which is not commonly observed. Therefore, it is of interest to gauge the conformation of the selectivity filter under those conditions using MD simulations. In open-gate E71V, Ba^2+^ stably binds to the selectivity filter at site S3 for 500 ns with Na^+^ (from the bulk solution) briefly binding to the outer sites of the selectivity filter (Figure 7c). The selectivity filter adopts a conductive state, but notably this conformation was not stable throughout the 500 ns simulation. The G77 Cα-Cα cross-subunit distances show that the filter cross-subunit distances fluctuate between 6 - 9 Å (Figure 7c, maximum near 400 ns) due to the conformational instability and deviation of the upper filter (Y78 to D80) in some subunits. Moreover, the selectivity filter is slightly narrower when compared with the canonical conductive filter (1K4C).^5^ For the closed-gate E71V with Ba^2+^ or K^+^ ions, the conformation of the selectivity filter is highly stable during 500 ns MD simulations. In both trajectories, Ba^2+^ or K^+^ ions predominantly bound at the same sites as the initial conformation (Figure 7 d, e). In comparison with the other simulations, the selectivity filter of the closed-gate E71V bound with Ba^2+^/K^+^ was very stable with a G77 Cα-Cα cross-subunit distance of ~ 8 Å, indicative of a canonical conductive filter.^5,61^

### W67 as a molecular switch

In the open-gate E71V crystal structures, the W67 sidechain is flipped 180° compared to WT KcsA, consistent with the ssNMR data (Figure 3b). To understand this conformational difference from a dynamical perspective, the orientation of W67 was monitored in all the trajectories. Based on the analysis, W67 in the open-gate E71V accessed multiple rotamer conformation (Figure 8). Starting from the equilibrated crystal structure and throughout the 500 ns simulation, the side chain of W67 in the open-gate E71V was very dynamic in nature. This dynamic behavior was observed in the simulations of open-gate E71V with K^+^, Na^+^ or Ba^2+^ (Figure 8a-c). Unlike the crystal structure, the conformation of W67 sidechain in the simulations of open-gate E71V was not confined to a certain conformation throughout the MD simulation. Rather, the side chain of W67 adopted multiple rotamer configurations. The high mobility of W67 in the open-gate E71V could be attributed to the orientation of W67 side chain. The 180° flipping of W67 side chain facing downward prevents its Nε1-Hε1 group from forming a hydrogen bond with the carboxylate group of D80, which is needed stabilize the configuration of W67 and D80, as well as for maintaining the structural integrity of the whole selectivity filter. In closed-gate E71V with Ba^2+^, the conformation of W67 in the crystal structure is identical to that in the WT KcsA of previous structures (1K4C, 1K4D). In the simulation, it is observed that the mobility of the W67 is significantly less than in the open-gate E71V simulations due to the stability of hydrogen bond between W67 and D80 (Figure 8d). The range of observed conformations of W67 in the crystal structures and ssNMR, indicates that this sidechain may access multiple rotameric states in the open-gate E71V. However, this is not the case in the closed-gate KcsA, presumably because the hydrogen bond restrains the mobility of W67.

**Figure 8.**
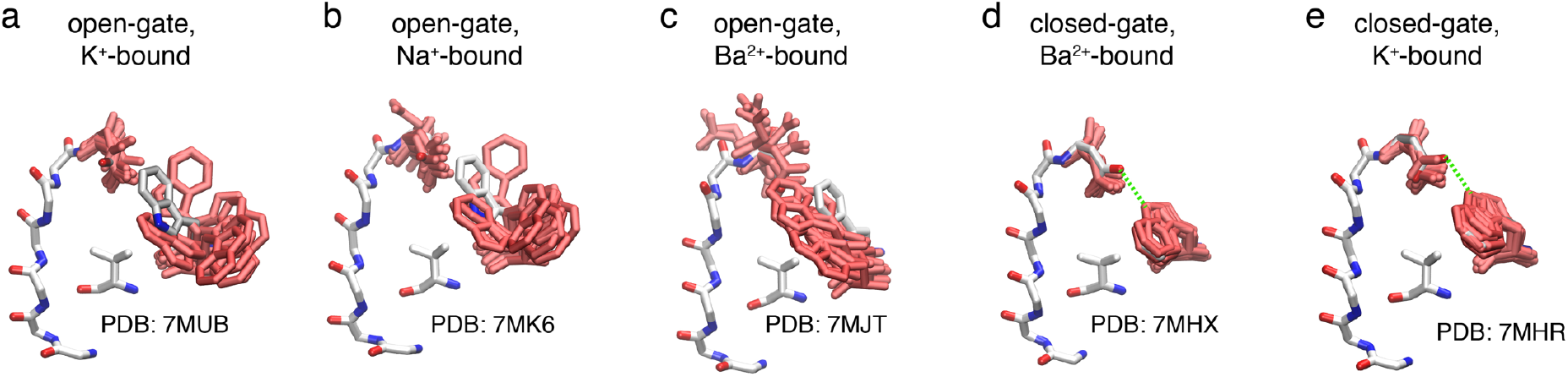
The dynamics of the W67 and D80 side chains in MD simulationsunder various conditions. (a) open-gate and K^+^ bound, (b) open-gate and Na^+^ bound, (c) open-gate and Ba^2+^ bound, (d) closed-gate and Ba^2+^ bound, and (e) closed-gate and K^+^ bound. The crystal structures are shown as reference (labeled with their PDB ID) in licorice representation and carbon atoms in white with multiple snapshots from simulations. The side-chain conformations are shown in red licorice, and these snapshots are selected for every 100 ns in all four subunits.

Unbiased MD simulations provide a powerful albeit limited way to explore the conformations accessible to a molecular structure; a 0.5 μs trajectory is likely to sample mainly the conformations around a meta-stable free energy basin. To more quantitatively assess the conformational preferences of the selectivity filter, free energy landscapes PMFs were calculated. The results shown in Figure 9 represents extensions of previous calculations.^29,31^ The computations are designed to allow a comparison of the WT and E71V mutant channels with fully open gate (~23 Å measured by Cα-Cα distance between T112). To obtain the free energy landscape, twodimensional potential of mean force (2D-PMFs) were performed using US/H-REMD simulations as a function of the Cα—Cα cross-subunit distance of G77 in the filter (coordinate r) and the position of the external K^+^ ion along the pore axis (coordinate z). To compare the propensity between the conductive conformation with K^+^ ion bound at S2 site and constricted structure with K^+^ at S1 site, two most typical occupancy observed in previous structures, the 2D PMFs only focus on the region covering the outer binding sites S1 and S2 (between 0 and 6 Å along the *z* coordinate).

**Figure 9.**
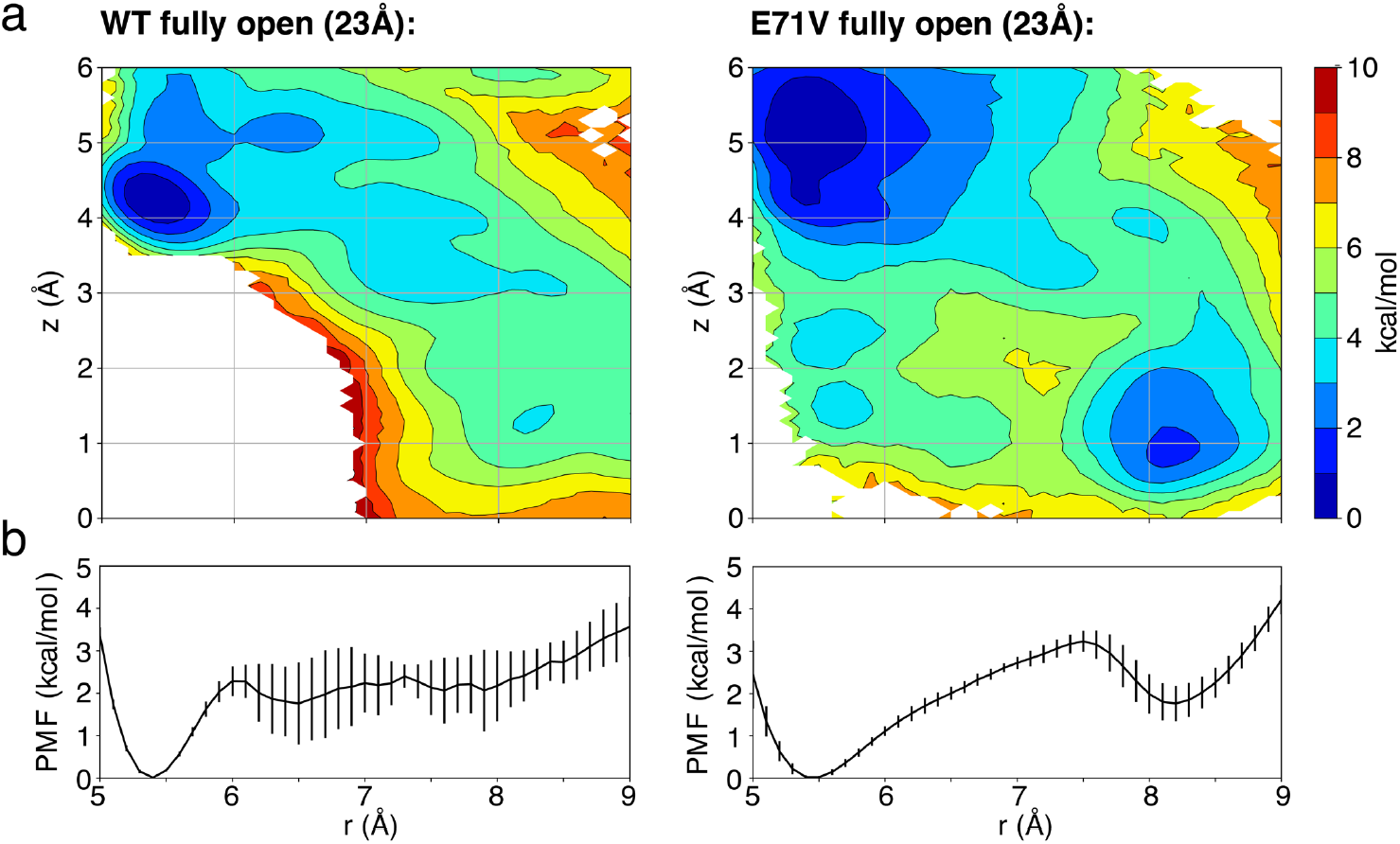
Free energy landscapes to assess the conformational preferences of the selectivity filter of WT and the E71V KcsA mutant as a function of intracellular gate opening. a) 2D PMF depending on two coordinates. The horizontal reaction coordinate r describes the width of the selectivity filter and is defined as the average cross-subunit pinching distance between the Cα atoms of G77. The vertical reaction coordinate z indicates the position of the external K^+^ ion along Z axis relative to the center of the selectivity filter. Extensions of previous calculations.^29,31^ b) 1D PMF projected along the horizontal reaction coordinate r (after Boltzmann integration of the vertical reaction coordinate *z*).

The calculated 2D PMFs clearly show the relative population differences for the conductive and constricted conformations between WT and E71V mutant (Figure 9a). For WT, there is deep free energy minimum around r = 5.4 Å corresponding to a stable constricted state, but no clear free energy basin in the conductive structural region (r ≈ 8.2 Å). It is reasonable to imagine that the selectivity filter would tend to spontaneously interconvert from the conductive to the constricted conformation for WT with an open intracellular gate, going downhill on the free energy surface (Figure 9b). For E71V mutant, there is clear minimum in the conductive region (r ≈ 8.2 Å), and the free energy difference is only 1.5 kcal/mol between the conductive and constricted minima. These features are broadly consistent with previous computational results obtained using a homology model of the pore domain the *Shaker* channel.^31^ The comparison of these two free energy landscapes (WT vs E71V) indicate that, although constricted conformation is favorable for both E71V and WT in open structure, there is higher probability to capture a conductive conformation for E71V mutant than WT. The presence of a single inactivating water at near the bottom of the filter frees the backbone of Y78 and G79 from hydrogen bonding constraints, which reduces the stability of the constricted state compared to the canonical constricted conformation of the WT (1K4D). This is consistent with a previous functional study showing that E71V is less inactivated than WT.^19^ It is also consistent with ssNMR data showing a shift of the V76CO signal in the conductive E71V filter (179.8 ^13^C ppm in E71V; 179.0 ^13^C ppm in WT KcsA) (Supplementary Figure 5), which, as we recently showed^44^, relates to a stabilization of the conductive filter. Interestingly, the free energy basin of constricted states is shallower and broader for E71V than WT, suggesting that the constricted state in E71V could represent a larger set of heterogeneous ensemble than WT.

## DISCUSSION

Using a combination of X-ray, ssNMR, and MD simulations the behavior of the E71V KcsA mutant was investigated. While the overall sequence identity between KcsA and the pore of mammalian Kv channels is on the order of 40%, the valine along the pore helix behind the filter is highly conserved among Kv channels suggesting that it is an important molecular determinant of function. X-ray crystallographic structures show that the filter of the open-gate E71V KcsA channel is narrowed but not constricted. This observation is in line with our ssNMR data, which also does not reveal the presence of the constricted filter conformation in inactivating conditions. In contrast with the canonical constricted state, the absence of bound water molecules behind the narrowed filter is noteworthy. Both X-ray and ssNMR unambiguously show the absence of the constricted state for the E71V KcsA whereas the MD simulations indicate that this conformation is possible. While a constricted state for the E71V filter is not captured by our experiments, it is difficult to rule out this possibility: a conformation extensively documented in the case of the WT KcsA should be physically accessible to the E71V mutant.

X-ray data with bound Ba^2+^ confirm that the substitution of E71 by a nonpolar valine alters the local selectivity of the cation binding sites and filter gating. The open-gate WT filter is constricted in the presence of Ba^2+^, which then either binds at the S4 site^42^ or, as shown by ssNMR, cannot bind at all to the filter. By contrast, the filter of open-gate E71V accommodates Ba^2+^ at the S3 site without constriction. The binding in the middle of the filter seems consistent with studies reporting that Ba^2+^ tightly binds to C-type inactivated *Shaker* and becomes trapped in its selectivity filter.^62,63^ In line with our data, Ba^2+^ binding in the middle of the filter was observed for the MthK^59^ and NAK2K^60^ channels that both have a valine at the position corresponding to E71 in KcsA. A structure of the MthK channel with constricted filter could never be captured,^37^ although this appears to be explained by differences in ion binding affinity.^38^

X-ray, ssNMR, and MD simulations show that the W67-D80 interaction behind the filter, critical for inactivation in Kv channels,^12,19,40,41^ breaks in E71V KcsA under inactivating conditions upon opening of the inner gate, consistent with the allosteric coupling between the activation gate and the selectivity filter.^21,22,29^ On the other hand, the W67-D80 interaction is intact in the constricted conformation of WT KcsA (1K4D), which aligns with the diverging role of W67: replacing W67 slows down inactivating in WT KcsA, whereas it promotes inactivation in Kv channels.^19^ Another interesting observation is that ssNMR data unambiguously show, based on Cαβ chemical shifts (Figure 3e), that the Y78 sidechain changes its conformation in E71V KcsA upon inactivation. Changes in the sidechain of F627 in hERG, equivalent to Y78 in KcsA, were suggested to modulate the inactivation mechanism^39,64^ and emulate the role of structural water molecules behind the filter. Consistent with this observation, we also recently showed that Y78 exhibits strongly enhanced dynamics if residue E71 in KcsA is replaced by an isoleucine,^44^ which is representative for channels like Kv2.1. Against this background, it generally seems that there is more to learn on the role the sidechains behind of the selectivity filter environment, given that the of substitution of a glutamic acid to a nonpolar valine at position 71 in the channel (Figure 6d) drastically increases the overall dynamics of the pore by a mechanism that we do not yet fully grasp.

Strikingly, ssNMR data show that the E71V selectivity filter can adopt two conformations under inactivating conditions. This could either mean the presence of two different symmetric conformations, or alternatively, a single asymmetric conformation. Non-symmetric filter conformations have been related to inactivation gating in the KcsA G77dA,^28^ TREK-1,^49^ and hERG channels.^39,58^ The MD trajectories of open-gate E71V KcsA display a filter that departs from 4-fold symmetry (Figure 7) and the calculated 2D-PMF shows a lower energy barrier in between constricted and conductive filter than in WT KcsA (Figure 9). Both of these factors could promote a more heterogeneous sampling of the conformational space in E71V. The heterogeneity of the inactivated E71V filter also aligns with a previous computational study^31^, and could be caused by the absence of inactivating water molecules behind the filter or by the rupture of the W67-D80 interaction. A potential reason why a second conformation in open-gate E71V KcsA was not observed by X-ray could be due to the presence of the stabilizing Fab fragments bound in the pore mouth vicinity could impair some of the natural conformational sampling,^18^ as recently shown by ssNMR.^44^

In pondering the different pieces of information, one should make clear that asking whether the E71V mutant can adopt or not the constricted filter conformation, and what is the true molecular mechanism of inactivation of this mutant are two distinct questions. With respect to the first question, the small magnitude of the energy difference between conductive and inactivated state in E71V KcsA must certainly be taken into consideration. The macroscopic current in E71V KcsA after a stimulus decays to about 57% from the peak current,^20^ corresponding to a free energy difference between the conductive and non-conductive functional states of about 0.17 kcal/mol. The ssNMR experiments are carried out with the full length channel, which inactivates less deeply than the truncated form used in for X-ray crystallography and MD simulations.^29^ The intracellular gate of the full-length opens only up to 19 Å whereas the truncated form used to calculate the 2D-PMF has a gate opening of 23 Å (Figure 9), which allosterically promotes inactivation (a smaller gate opening would allosterically shift the equilibrium toward the conductive state). Moreover, the channel crystals include the presence of Fab antibodies that destabilize the inactivated state.^18^ With respect to the second question, it seems likely that there exists some non-constricted state of the E71V filter that is functionally non-conductive—a state that is associated with the functional phenotype of inactivation observed in the electrophysiological experiments. Strengthening this interpretation is the fact that the sample probed by ssNMR, with a full-length KcsA E71V channel incorporated into a fluid membrane, reflects faithfully the conditions imposed by the functional electrophysiological measurements (apart from the non-equilibrium ion current that perhaps promotes entry into C-type inactivation).

Choosing to wander on the side of caution, we conclude that there seems to exist at least two conformational states of C-type inactivation in the case of the Kv-like E71V mutation of KcsA. There is the familiar conformation in which the filter is constricted at the level of the central glycine, and there is a second conformation in which the filter is not constricted but slightly narrowed over its entire length. Such a non-conductive and non-constricted conformation was recently buttressed by some experimental studies with Kv channels.^13,35–38^ While it is generally non-trivial to imagine how a non-constricted filter could act to robustly inhibit K^+^ current, the concept has been discussed previously by Nimigean, Bernèche, and co-workers.^36^ Although this scenario is not captured by MD simulations (the narrowed pore can rapidly relaxes in the presence of K^+^), the concept of a non-conductive narrowed filter in the Kv-like channel is supported here by a transition from a highly dynamic conductive filter to very rigid filter under inactivating conditions, as indicated by ssNMR. Like in a bench vise, the four rigidly and compactly packed E71V protomers would block the passage of K^+^ through the filter. Coupling between pore helix and transmembrane helices could modulate the packing of the filter.^36^ One may speculate that the presence of the voltage-sensor could further affect the packing of the pore-domain. Such a narrowed-rigid state could possibly correspond to a pre-inactivated functional state that could ultimately interconvert into the long-lived constricted state.

## METHODS

### Crystallization and structure determination

The open-gate E71V KcsA was expressed as reported in Cuello et al., 2017.^23^ The pQE32 expression vector containing the coding sequence for E71V closed-gate KcsA channel was transformed into *E. coli* strain XL-1 Blue cells. Fresh LB media was inoculated from an overnight culture and grown at 37°C with shaking until reaching an OD_600_ of 0.8-1.0, after which expression was induced by adding 0.5 mM isopropyl-β-D-thiogalactopyranoside (IPTG). Growth was continued for 3 hours at 37°C. After this step the open- and closed-gate KcsA were treated similarly. Cell pellets were resuspended in lysis buffer (20 mM Tris, pH 8.0; 0.2 M NaCl) supplemented with 1mM PMSF and 20 μg/ml of DNase I. The resuspended cells were lysed by homogenization using an Avestin Emulsiflex C5. The cell membranes were collected by centrifugation at 100,000 xg at 4°C for 1 hour. The membrane pellet was resuspended in 20 mM Tris, pH 7.5, 0.15 M KCl and solubilized by incubation with 40 mM DM (n-Decyl-β-D-maltoside) overnight at 4°C. The sample was centrifuged at 100,00 xg for 1 hour and the supernatant was loaded on a HisTrap FF 5ml nickel metal affinity column previously equilibrated with buffer A (20 mM Tris, pH 7.5, 0.15 M KCl, 5 mM DM, 20 mM imidazole) and eluted with buffer B (buffer A + 300 mM imidazole). The C-terminal tail was removed by incubation with 1 μg/ml chymotrypsin for 1 hour at room temperature. The chymotrypsin-treated protein was further purified by size exclusion chromatography using a Superdex 200 HR 10/30 column with an AKTA FPLC system (GE Healthcare) in buffer C (50 mM Tris, pH 7.5, 0.15 M KCl, 5 mM DM). The peak fraction was collected and incubated with KcsA-Fab and size exclusion chromatography was repeated to remove excess Fabs. To obtain KcsA under Ba^2+^ conditions, the purified KcsA-Fab complex was incubated with 5 mM BaCl2 prior to crystallization and the KCl is replaced by 0.15 M NaCl in the final size exclusion chromatography buffer.

The KcsA-Fab complex was concentrated to 15 mg/ml and crystallized in 50 mM magnesium acetate, 50 mM sodium acetate pH 5.5 and 25% PEG 400 at 20°C. After 5 days, crystals were harvested and soaked in cryoprotectant solution containing 40% PEG 400 for 1 minute at room temperature. This step was repeated 3 times followed by flash-freezing in liquid nitrogen. X-ray diffraction datasets were collected at the NECAT 24-ID-C beamline at the Advanced Photon Source. Crystal structures were determined by molecular replacement using the structures 1K4C and 6W0A as search models. The anomalous difference Fourier maps were calculated using Phenix. The structure refinements were performed using REFMAC. The electron density map and KcsA models were inspected by COOT.

### Solid-State NMR spectroscopy

Inversely Fractionally deuterated^43^ WT KcsA and E71V mutant channels were expressed in *E. coli* M15 cells (Qiagen) using H_2_O-based M9 medium supplemented with ^15^NH_4_Cl and D-glucose-^13^C6-d7. Cells were subsequently harvested, treated with lysozyme, and lysed via French press. The membranes containing the channels were collected by centrifugation (100,000×*g*) and proteins were extracted with 40 mM DM.^65^ Channels were purified using Ni-NTA agarose beads (Qiagen). Liposome reconstitution was performed using *E. coli* polar lipids (Avanti) at 1:100 protein:lipid molar ratio co-solubilized in DM, in which the detergent was removed using polystyrene beads (Bio-Beads SM-2).^65^ Before the ssNMR measurements, reconstituted samples were suspended in phosphate buffer (pH 7.4, 100 mMK^+^) or citrate buffer (pH 3-4, 0 mM K^+^). Ba^2+^-binding studies were conducted with HEPES buffer (pH 7, 5 mM BaCl2) and citrate buffer (pH 4, 5 mM BaCl2) to probe closed-gate and open-gate states, respectively. Samples were incubated in the buffers for 2 days.

3D ssNMR experiments^44^ (CANH, CONH, CAcoNH, COcaNH) for sequential backbone assignments were performed at 800 MHz (^1^H-frequency) using 60 kHz magic angle spinning (MAS) frequency and a real temperature of approximately 305 K. See Supporting Information for a list of 3D experiments. 2D ^13^C–^13^C PARISxy^66^ experiments for side chain chemical shift assignments were performed at 700 MHz using 42 kHz MAS and a ^13^C–^13^C mixing time of 110 ms. ^15^N *T*_1rho_ relaxation experiments were run at 700 MHz and 60 kHz MAS using a ^15^N spin lock amplitude of 17.5 kHz.^44^ PISSARRO^67^ decoupling was used as decoupling method in all dimensions. 2D ^15^N(^1^H)^1^H ssNMR experiments^48^ using 1.5 ms ^1^H^1^H spin diffusion water was used to probe the presence of structural water behind the filter.

### Molecular dynamics simulations

The open and closed-gate crystal structures of the KcsA E71V mutant obtained in the present study were used as the starting model for MD simulations. The channel was embedded in a bilayer of 3POPC:1POPG lipids and solvated in 150 mM NaCl using the web service CHARMM-GUI.^68,69^ Most residues were assigned their standard protonation state at pH 7. The total number of atoms in the atomic model is on the order of 41,000. The all-atom CHARMM force field PARAM36 for protein,^70–73^ lipids,^74^ and ions^75^ was used together with the TIP3P water model.^76^ The atomic channel models were refined using energy minimization for at least 2,000 steps, and the ions and backbone atoms (except those from the selectivity filter) were kept fixed throughout the minimization procedure. All the simulations were performed under NPT (constant number of particle N, pressure P, and temperature T) conditions at 310 K and 1 atm, and periodic boundary conditions with electrostatic interactions were treated by the particle mesh Ewald method^77,78^ and a real-space cutoff of 12 Å. The simulations use a time step of 2 fs, with bond distances involving hydrogen atoms fixed using the SHAKE algorithm.^79^ After minimization, positional restraints on all the Cα atoms were gradually released after which a trajectory of 500 ns was generated using NAMD version 2.11.^80,81^ The biased simulations to calculate the two-dimensional (2D) potential of mean force (PMF) surfaces were generated with Hamiltonian replica-exchange MD (US/H-REMD) ^82^ using the multiple-copy facility of NAMD^83^ and were unbiased using the weighted histogram analysis method (WHAM).^84,85^ Analysis was carried out using the program VMD.^86^

## Data availability

Data supporting the findings of this manuscript are available from the corresponding author on reasonable request.

## Author Contributions

AR did X-ray crystallography studies; AR, JL, and BR performed MD simulations; LB did molecular biology; BJAV, FK, FN, JMS, and MW acquired ssNMR data. All authors contributed to the data analysis and drafting of the manuscript.

## Acknowledgments

This work was funded in part by the National Institutes of Health/National Institute of General Medical Sciences (NIH/NIGMS) through grants R01-GM062342 and U54-GM087519, and the Dutch Research Council (NWO, grant numbers 723.014.003 and 711.018.001).

