## Supplementary Information for "A different mechanism of C-type inactivation in the Kv-like KcsA mutant E71V"

| | E71V closed-gate | | E71V open-gate | | |
|---|---|---|---|---|---|
| | $K^+$ | $Ba^{2+}$ | $Na^+$ | $Ba^{2+}$ | $K^+$ |
| | 7MHR | 7MHX | 7MK6 | 7MJT | 7MUB |
| Wavelength (Å) | 0.9792 | 0.9792 | 0.9792 | 0.9792 | 0.9792 |
| Source | APS 24-ID-E | APS 24-ID-E | APS 24-ID-E | APS 24-ID-E | APS 24-ID-E |
| Resolution (Å) | 2.7 | 2.9 | 3.1 | 3.3 | 3 |
| Space group | I 4 | I 4 | I 4 | I 4 | I4 |
| Cell parameters (Å) | 155.8, 155.8, 76.2 | 155.2, 155.2, 75.3 | 140.1, 140.1, 69.5 | 140.5, 140.5, 69.7 | 139.5, 139.5, 69.2 |
| (°) | 90.0, 90.0, 90.0 | 90.0, 90.0, 90.0 | 90.0, 90.0, 90.0 | 90.0, 90.0, 90.0 | 90.0, 90.0, 90.0 |
| Total reflections | 155,982 | 126,948 | 79,887 | 62,362 | 87679 |
| Unique reflections | 23,491 | 21,076 | 12,391 | 10,372 | 13389 |
| Multiplicity | 6.6 (6.6) | 6.0 (6.0) | 6.4 (5.9) | 6.0 (6.0) | 6.5 (6.4) |
| Completeness (%) | 99.7 (98.1) | 99.8 (100.0) | 99.9 (99.9) | 99.9 (100.0) | 99.5 (99.8) |
| Mean I/sigma(I) | 10.3 (2.9) | 9.0 (0.7) | 8.3 (0.6) | 11.3 (1.5) | 12.5 (1.0) |
| R-merge | 0.20 (1.03) | 0.13 (2.7) | 0.09 (2.5) | 0.06 (1.18) | 0.061 (1.87) |
| R-pim | 0.12 (0.64) | 0.09 (1.78) | 0.05 (1.71) | 0.04 (0.77) | 0.038 (1.21) |
| CC1/2 | 0.97 (0.88) | 0.92 (0.14) | 0.99 (0.16) | 0.99 (0.65) | 0.999 (0.35) |
| Reflections used in refinement | 22,301 | 17,580 | 11,693 | 9,809 | 12,659 |
| R-work/R-free | 0.17/0.20 | 0.20/0.24 | 0.21/0.25 | 0.21/0.26 | 0.19/0.25 |
| RMS bond length (Å) / angle (°) | 0.01/2.1 | 0.01/1.9 | 0.01/1.8 | 0.01/1.6 | 0.01/1.8 |
| Ramachandran favored (%) | 91.6 | 89.6 | 78.5 | 79.6 | 78.2 |
| allowed (%) | 7.7 | 9.7 | 20.3 | 19.2 | 20.6 |
| outliers (%) | 0.7 | 0.7 | 1.2 | 1.2 | 1.2 |

*values in parentheses are for highest-resolution shell.

**Table 1.** Data collection and refinement statistics for open and closed gate E71V KcsA.

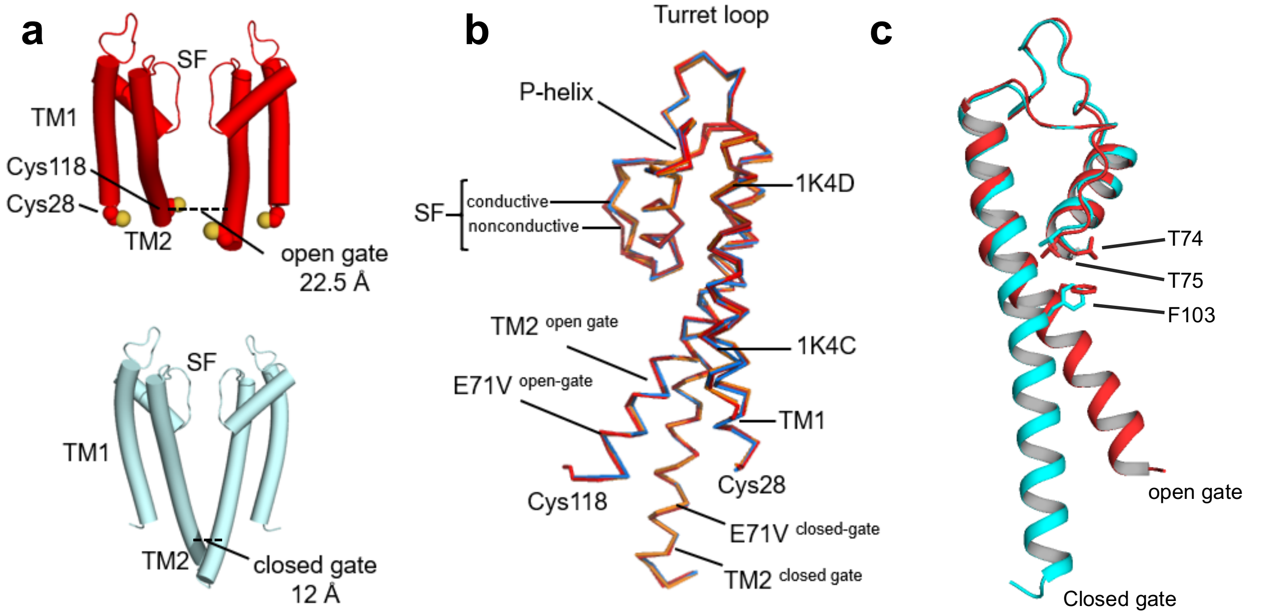


**Supplementary Figure 1.**

**X-ray structures of E71V open and closed gated KcsA in an allosteric network.** a) Tube representation of open E71V KcsA (red) and closed (cyan) WT KcsA. Two monomers are shown for clarity showing the transmembrane helices (TM) and selectivity filter (SF). The cysteine residues (28 and 118) used to produce a lock-open KcsA are depicted in red and yellow spheres. The T112-T112 distance is shown in dashed lines. b) Superposition of open-gate E71V KcsA and closed-gate KcsA with crystal structures 1K4C and 1K4D. One monomer is shown in ribbon representation. c) Superposition of X-ray structures of closed-gate and open-gate KcsA (rotated 180$^{o}$ with respect to figure b, for clarity) showing the F103 sidechain changes rotamer upon channel opening, moving closer to the T74 and T75 sidechains. The interaction between F103 and T74/T75 an important determinant of the gate-filter allosteric coupling in KcsA[33].

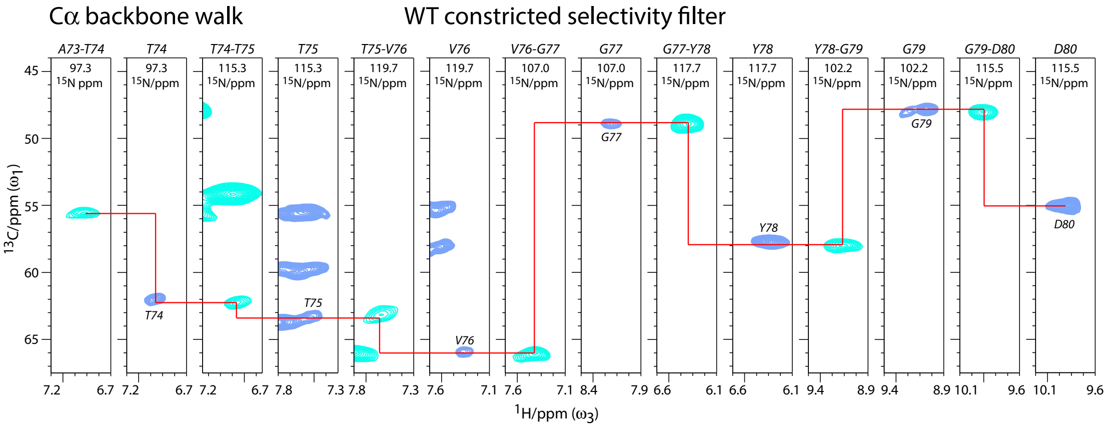


### Supplementary Figure 2.

**Sequential ssNMR assignments with $^{1}$H-detected 3D experiments.** Cα-Cα$^{+1}$ backbone walk showing full connectivity for the entire constricted selectivity filter (A73 – D80) in WT KcsA, acquired under acidic conditions (pH4, 0 mM K$^{+}$). Dark blue signals show CAH slices from a 3D CANH experiment, cyan CAH slices were taken from a 3D CAcoNH experiment.

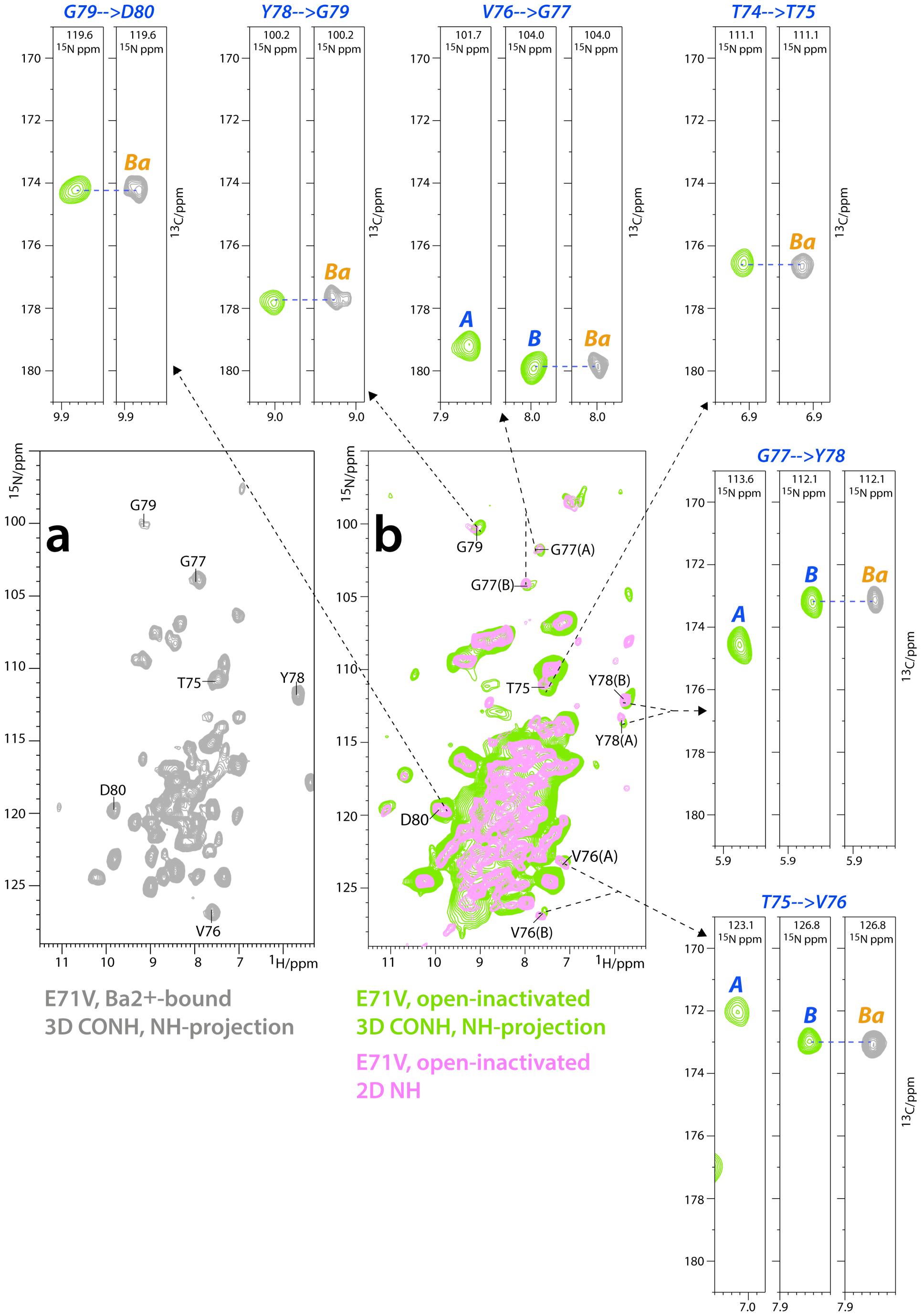


**Supplementary Figure 3**

**NMR assignments of the two conformations of the inactivated E71V selectivity filter.** a) 2D NH projection from a 3D CONH spectrum (in grey) of $Ba^{2+}$-bound E71V (pH7, 5 mM $Ba^{2+}$). Selectivity filter signals are annotated. b) Superposition of 2D NH projection from a 3D CONH spectrum (in green) and a 2D NH spectrum (magenta) of open-inactivated E71V (pH3, 0 mM $K^{+}$). The two conformations *A* and *B* of the inactivated filter are labelled in the spectrum. $^{15}$N-$^{1}$H strips form the 3D CONH experiments are shown for all selectivity filter signals T74-G79, which shows that conformation *B* is similar to the $Ba^{2+}$ conformation.

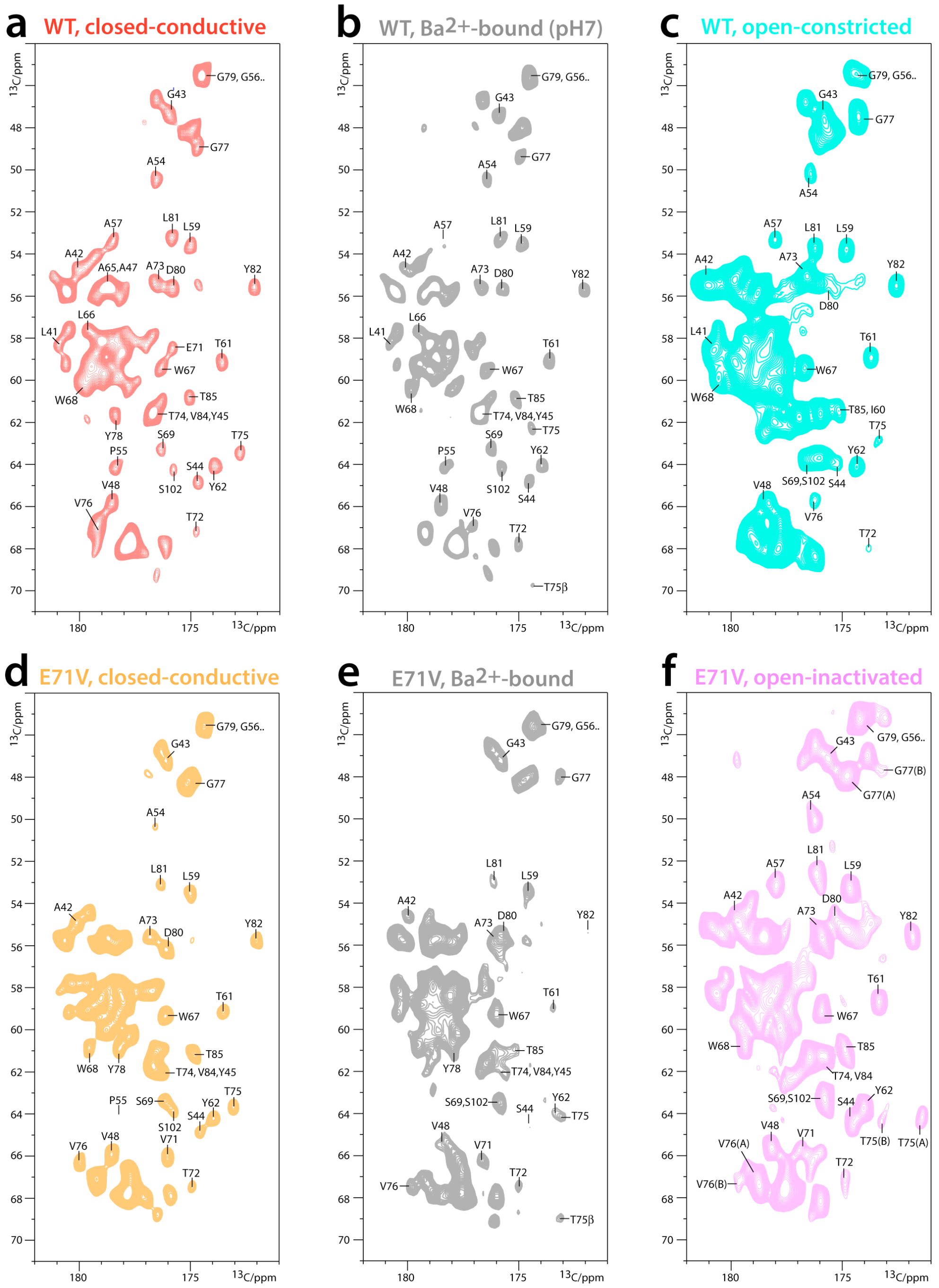


**Supplementary Figure 4**

**2D CC ssNMR spectra of WT and the E71V KcsA in different states.** 2D ssNMR PARIS $^{13}C^{13}C$ spectra acquired at 700 MHz and 42 kHz MAS. a) WT KcsA closed-conductive (pH7, 100 mM $K^+$). b) WT $Ba^{2+}$-bound (pH7, 5mM $Ba^{2+}$). c) WT open-constricted (pH4, 0 mM $K^+$), d) E71V KcsA closed-conductive. e) E71V $Ba^{2+}$-bound (pH7, 5mM $Ba^{2+}$). f) E71V open-inactivated (pH3, 0 mM $K^+$).

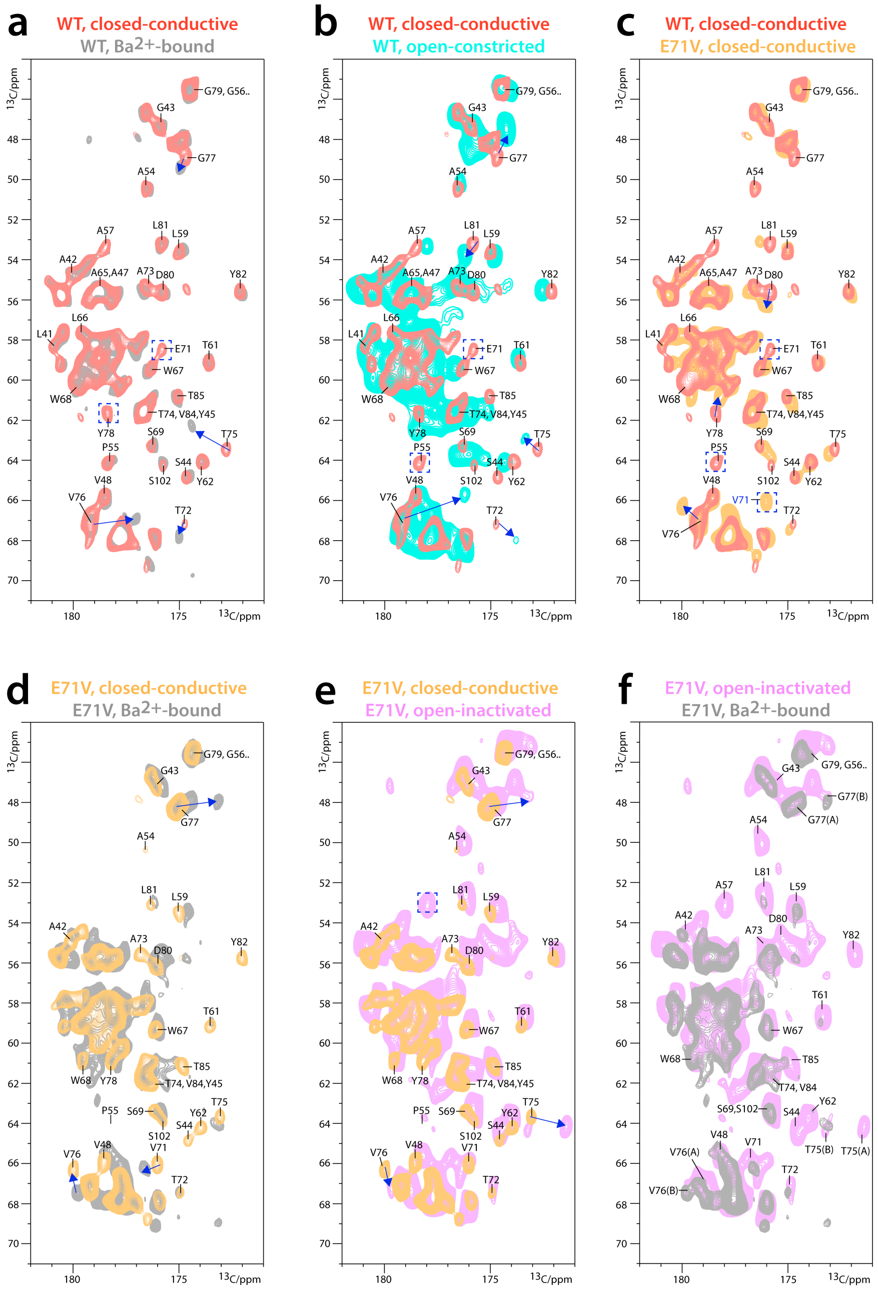


### Supplementary Figure 5

**Comparison of 2D CC ssNMR spectra.** Superposition of 2D ssNMR PARIS $^{13}C^{13}C$ spectra acquired at 700 MHz and 42 kHz MAS. a) WT KcsA closed-conductive (pH7, 100 mM $K^+$) on WT $Ba^{2+}$-bound (pH7, 5mM $Ba^{2+}$). b) WT KcsA closed-conductive on WT open-constricted (pH4, 0 mM $K^+$). c) WT KcsA closed-conductive on E71V closed-conductive. d) E71V closed-conductive on E71V $Ba^{2+}$-bound (pH7, 5mM $Ba^{2+}$). e) E71V closed-conductive on E71V open-inactivated (pH3, 0 mM $K^+$). f) E71V $Ba^{2+}$-bound on E71V open-inactivated.

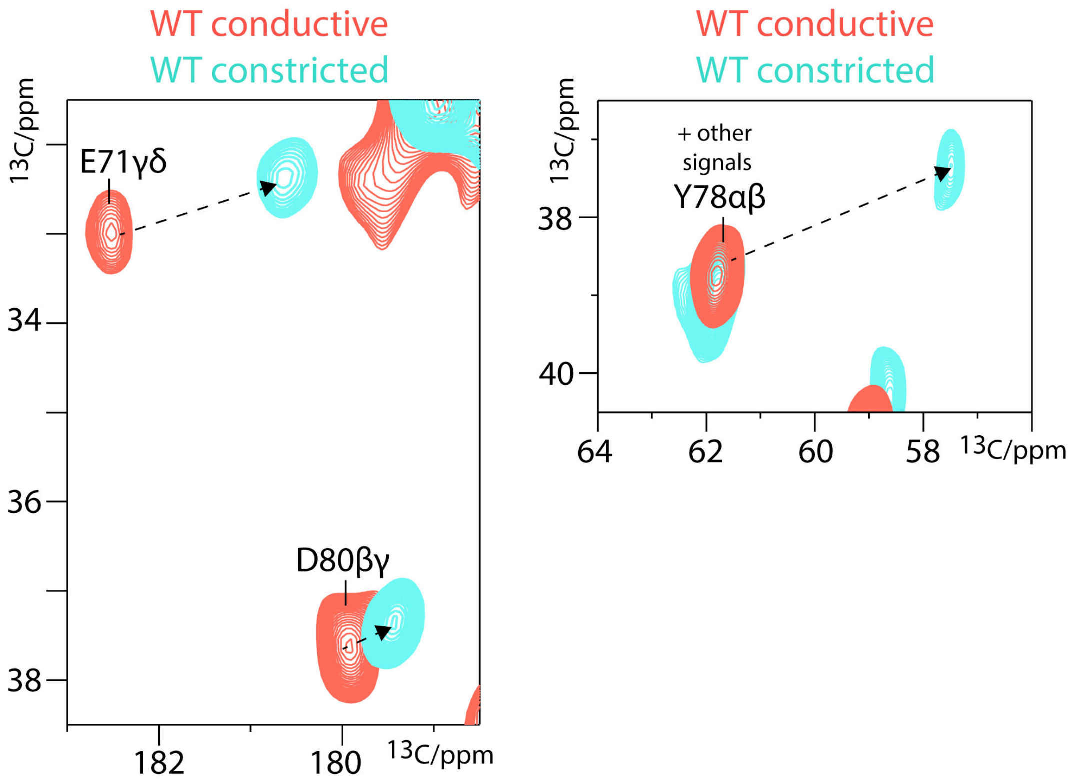


**Supplementary Figure 6**

**ssNMR shows major conformational changes in the hydrogen bonding network behind the filter of WT KcsA upon inactivation.** A comparison of 2D CC PARIS ssNMR spectra acquired with WT KcsA in closed-conductive (red) and open-constricted (cyan) states shows strong chemical shift changes in the sidechain of E71 and the backbone of Y78. This is consistent which is break of the hydrogen bond between E71 – Y78, as shown in the 1K4D crystal structure.

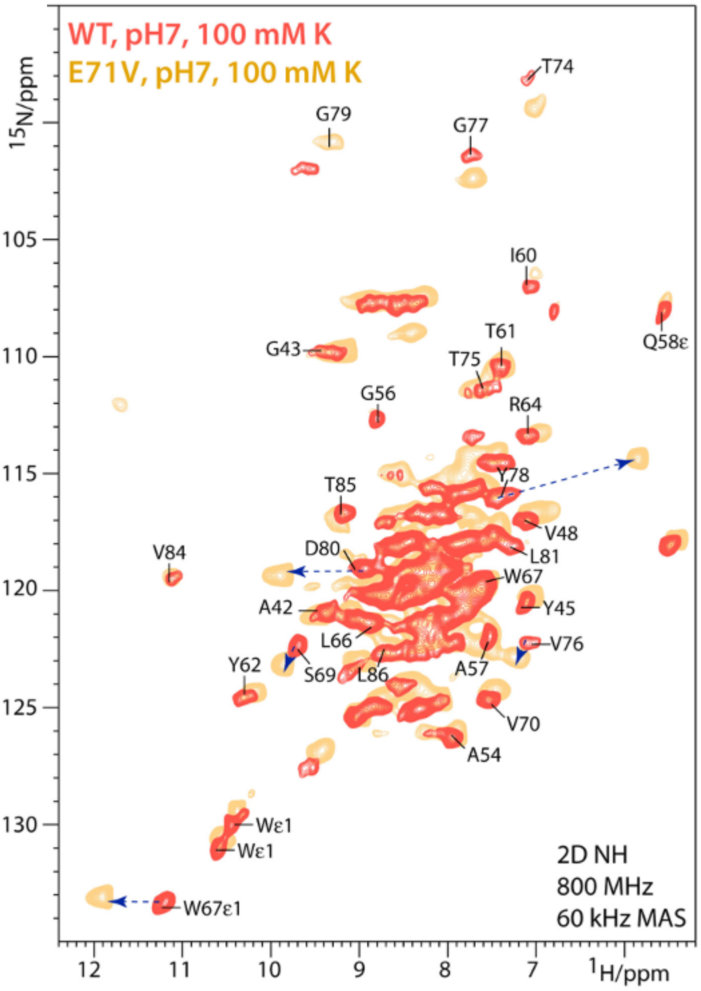


**Supplementary Figure 7**

**ssNMR comparison of closed-conductive E71V and WT KcsA.** Comparison of 2D NH spectra of the closed-conductive states of WT (red) and E71V KcsA (orange). Next to changes in the selectivity filter, we observe major CSPs for the sidechain of W67, and the backbone of Y78 and D80. These chemical shift changes are, presumably, due to the non-polar nature of the V71 sidechain, which cannot form hydrogen bonds with Y78 and D80.

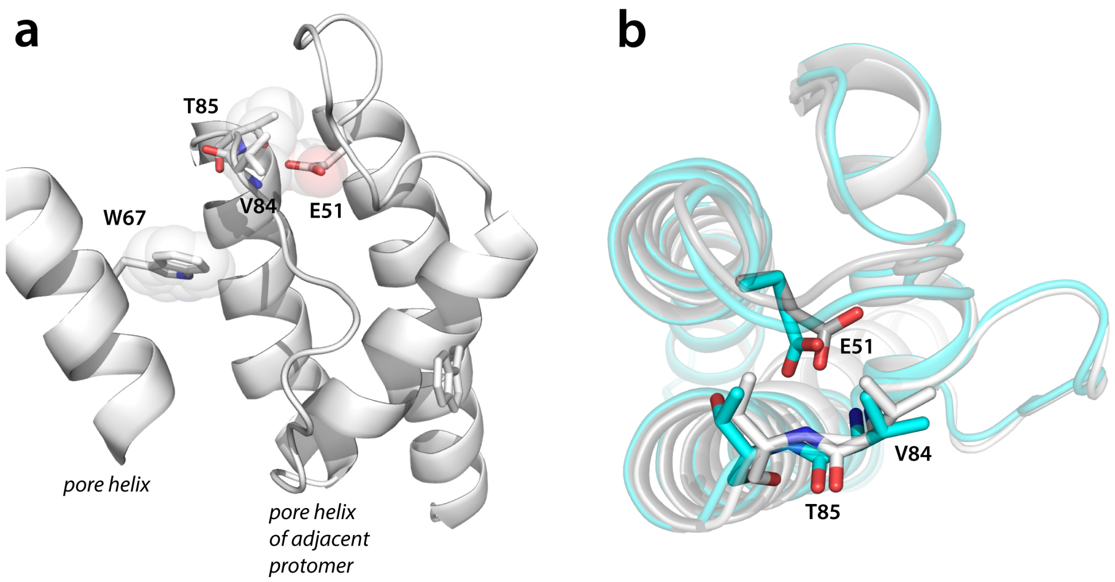


**Supplementary Figure 8**

**Allosterically induced rotameric change of W67 propagates to the turret region.** a,b) X-ray and ssNMR data show that the opening of the inner gate allosterically modulates the conformation of W67 of the pore helix. This propagates to the distal turret residues V84 and T85 in the adjacent protomer, right above the W67 sidechain, that form a critical hydrogen bond with the sidechain of E51.
